# Sign language communication enhances representations for hands in high-level visual cortex

**DOI:** 10.64898/2026.08.14.744668

**Authors:** Larissa Kahler, Klaudia Grote, Merve Sarac, Klaus Willmes, Kerstin Konrad, Marisa Nordt

**Affiliations:** AG Developmental Cognitive Neuroscience, Institute of Medical Psychology and Medical Sociology, University Hospital RWTH Aachen, Pauwelsstraße 19, 52074 Aachen, Germany; Competence Center for Sign Language and Gesture (SignGes) at RWTH Aachen University, Theaterplatz 14, 52062 Aachen, Germany; Institute of Medical Psychology and Medical Sociology, University Hospital RWTH Aachen, Pauwelsstraße 19, 52074 Aachen, Germany; Clinic for Neurology, University Hospital RWTH Aachen, Pauwelsstrasse 30, 52074 Aachen, Germany; Forschungszentrum Jülich GmbH (INM-10), Wilhelm-Johnen-Straße, 52428 Juelich, Germany

## Abstract

High-level visual cortex supports the perception and recognition of visual categories. What is the role of experience in shaping this region and the time course over which it stays malleable? We tested whether experience with a sign language, where information is conveyed via the hands and face, shapes category representations in high-level visual cortex. We acquired functional MRI data from 20 hearing signers and 20 non-signers while they viewed images from ten categories, including faces and hands. We compared the neural distinctiveness and size of category-selective regions between groups in ventral temporal and lateral occipito-temporal cortex. Signers showed higher distinctiveness for hands and larger hand-selective regions than non-signers in the left hemisphere, and sign language experience predicted neural hand representations. Critically, ventral hand representations were also enhanced in signers who learned sign language in adulthood. These findings indicate that visual cortex retains experience-dependent plasticity into adulthood, with implications for visual learning and cortical development.

## Introduction

The recognition and perception of visual categories such as faces, body parts or written words is essential for everyday life. Human high-level visual cortex, including the ventral temporal cortex (VTC) and the lateral occipitotemporal cortex (LOTC), is critical for these processes^1–3^. In fact, high-level visual cortex contains functional regions that show a selective response to categories such as faces^4–6^, body parts^1,7–9^, or written words^10^. Importantly, while category-selective regions are causally involved in the perception of categories^11–15^, information on these categories is also represented in distinct and reproducible distributed response patterns across high-level visual cortex^16,17^. Central questions concern the extent to which experience shapes this functional organization, and the time window over which it remains malleable.

Prior research focusing on reading has enhanced our knowledge about effects of experience on high-level visual cortex. These studies revealed that extensive experience with written words during reading acquisition - typically in childhood - leads to the formation of word-selective regions^10,18–21^. Critically, the size of these regions is linked to reading ability^22^ and their response profile is tuned to the learned script^23^. In addition, only adults who have learned to read, but not illiterates, show an enhanced response to script compared to other visual stimuli like faces in the location of word-selective regions^24^. Beyond the development within these focal regions, learning to read also shapes distributed responses to words over ventral temporal cortex^25,26^.

While these results demonstrate how experience shapes word-selective regions, important questions regarding the experience-dependent plasticity of high-level visual cortex remain. First, are other category-selective regions similarly shaped by extensive experience with their respective categories? Notably, word-selective regions differ from other category-selective regions in several aspects that are particularly relevant to their malleability. Word-selective regions are the only category-selective regions in VTC emerging only with the onset of schooling^19–21,27^, when regions selective to other visual categories, such as faces or bodies, have already emerged^28–30^. Further, and possibly related to their late emergence, word-selective regions are more variable regarding their location compared to other category-selective regions^31,32^. Critically, prior results show that during their emergence, word-selective regions recycle^33^ cortex previously involved in the perception of limbs and hands^34,35^, suggesting a particularly high degree of malleability of this part of cortex during childhood.

In addition, for other categories such as faces or places, it is more difficult to link visual experience to neural development, because (i) extensive exposure to these categories begins at birth and lacks the clearly defined onset that literacy acquisition provides, and (ii) unlike reading experience, the kind and extent of everyday experience with these categories is difficult to capture. For instance, limb-selective regions shrink from childhood to adulthood^35,36^ and limb- selectivity gets recycled into word-selectivity^35^, but the reason for the shrinking of limb-selective regions is currently unknown. A current hypothesis links the shrinking of limb-selective regions to changes in children’s viewing behavior^34,35^, as research using eye-tracking has shown that children look more often and longer at hands compared to adults^37^. This link, however, has yet to be tested.

A second important question is to what degree experience can shape distributed responses and category-selective regions beyond childhood, i.e., in the adult brain. Prior evidence on this question is mixed. On the one hand work from animal studies suggests strong limits to the plasticity of high-level visual cortex in the adult compared to the juvenile brain. Research in macaques undergoing symbol training suggests that while juvenile monkeys developed a cortical representation for the learned symbols in the temporal lobe, adult monkeys did not^38^. On the other hand, different lines of evidence suggest that high-level visual cortex retains at least some degree of plasticity into adulthood. For instance, studies on categorization and discrimination training in human adults suggest that category training enhances voxel-selectivity^39,40^, though not in a consistent anatomical location across participants. Moreover, face-selective regions, continue to mature throughout adolescence^41^ and at least until age 17^35^ – a particularly prolonged developmental trajectory. Although maturation cannot be excluded as an explanation for this development, it is plausible that these findings reflect ongoing experience-dependent plasticity, possibly driven by the expanding social circles and increasing face exposure during adolescence.

To address these gaps in knowledge, this study tests whether communication with a sign language, where information is conveyed visually through movements of the face and hands, shapes high-level visual regions. In fact, sign languages are known to sculpt the visual system and specific visual and visuospatial skills^42^, e.g. enhance mental rotation skills^43^ and enlarge the lower visual field^44^. Further, signers perform more accurately on face identity recognition tasks^45,46^ (but see^47^) and in the recognition of some facial expressions^48^ compared to non-signers. Behavioral results also indicate that the representation of hand size is altered in signers compared to non- signers^49^.

In combination, these behavioral results raise the question whether communication with sign languages shapes the neural representations for faces and hands, and if so, whether this occurs exclusively before the onset of adulthood. Prior evidence suggests different hypotheses: One hypothesis is that communication with sign languages shapes representations for faces and hands because prior literature suggests that high level visual regions are influenced by extensive visual experience. In addition to the effects of learning to read on high-level visual cortex outlined above^10,18–22,24–26,50^, extensive visual experience during development shapes VTC organization more broadly: playing Pokémon during childhood results in a distinct representation for Pokémon in VTC^51^ and protheses use is associated with more stable response pattern for prostheses in the lateral body-selective region compared to non-users^52^. These results suggest that extensive visual experience shapes both distributed responses and the size of category-selective regions in high-level visual cortex.

On the other hand, it is also possible that communication with a sign language does not have an effect on category representations in high-level visual cortex because (i) hands are mostly perceived in the periphery during communication with sign language as signers fixate mostly on the face^44,53^, and (ii) faces are also typically fixated on in communication with spoken language^54^.

A third hypothesis is that sign language use shapes the representation for faces and hands but only if sign language was acquired before adulthood. This hypothesis is supported by prior research suggesting that literacy acquisition affects the visual system differently depending on whether reading was learned in childhood or adulthood^24,50^, and results showing reduced plasticity in adult-onset blind individuals compared to congenitally blind individuals^55^, both implying that the adult visual system is less susceptible to experience-dependent reorganization.

To differentiate between these hypotheses, we compared category representations for faces and hands in high-level visual cortex in 20 participants with knowledge of German Sign Language (signers) and 20 participants without sign language knowledge (non-signers). Participants in both groups had no hearing impairments and were matched for age (non-signers: mean age= 35.1y, SD = 11.59; signers: mean age= 34.8y, SD= 13.06, BF_10_ = 0.32, δ = -0,06, 95% CrI [-0.62, 0.49]).

Critically, signers communicated in sign language extensively with a mean of 20 hours per week (SD=14.65; **Fig. 1A**). The average age of sign language acquisition in our sample was 17 years (M =17.7 years, SD=8.33), with the majority of our sample (16 of 20) having acquired sign language as adults and only 4 before the onset of adulthood (3 from birth and one during adolescence).

**Figure 1.**
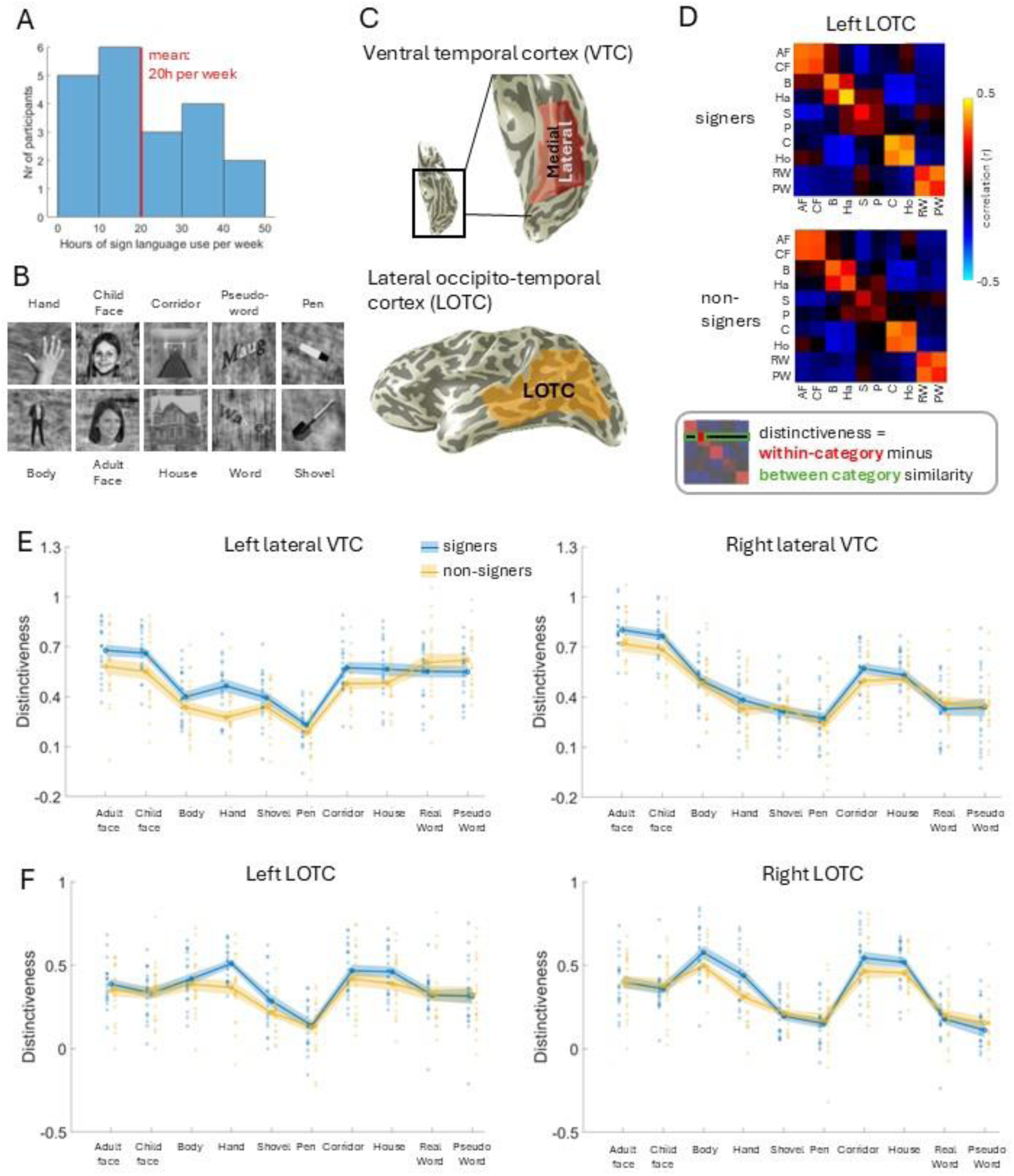
Communication with a sign language enhances distributed responses to hands in high-level visual cortex. **A** Histogram of hours per week spent using sign language to communicate, among signers (N=20). The average sign language use is 20 hours per week. **B** Example stimuli for the ten categories shown during fMR imaging. The 10 categories can begrouped into 5 broader domains (bodies, faces, places, words, tools). The faces illustrated were not presented in the real experiment and are the faces of two of the authors (M.N. and L.K), one of them as a child. **C** Examples for anatomical regions of interest (ROIs), which were defined on each participant’s individual brain surface based on anatomical landmarks: the medial and lateral ventral temporal cortex (VTC, top) and the lateral occipito-temporal cortex (LOTC. bottom). **D** Mean Representational Similarity Matrices (RSMs) for signers (top, N=20) and non-signers (bottom, N=20) in the left LOTC ROI. AF= adult faces, CF=child faces, B=bodies, Ha=hands, S=shovels. P=pens. C=corridors, Ho=houses,RW=re3l words. PW=pseu do words. Box: Illustration of the computation of category distinctiveness based on an RSM. **E, F** Distinctiveness for the ten categories in the left and right lateral VTC (panel E) and in the left and right LOTC (panel F) for signers (blue, N=20) and non-signers (yellow. N=20). Unfilled circles* mean, shaded areas = standard error of the mean, filled circles* individual distinctiveness values.

Participants took part in structural and functional magnetic resonance imaging (MRI). During functional imaging, participants completed three runs of an experiment^56^ in which they watched images of ten categories (**Fig. 1B**) including images of faces and hands (some also showing arms) while performing an oddball task. After scanning, participants completed questionnaires on handedness and sign language acquisition and use.

To assess differences in category representations in high-level visual cortex between the groups, we first defined three anatomical regions of interest (ROIs) per hemisphere on each participant’s individual inflated cortical surface based on cortical landmarks. To capture category representations on the ventral surface of the temporal lobe, we manually delineated VTC ROIs, which were separated into a medial and a lateral part (**Fig. 1C -top)**. To capture category representations on the lateral surface, we manually delineated LOTC ROIs (**Fig. 1C -bottom**; for definition criteria see section Methods). We next assessed both the distributed response patterns and the number of category-selective voxels for the 10 categories in these ROIs. In addition, we also manually delineated face- and hand-selective ROIs in each participant.

## Results

### Higher distinctiveness for hands in high-level visual cortex of signers compared to non-signers

We first asked if communication with German Sign Language shapes distributed response patterns to hands and faces in VTC and LOTC. To answer this question, we determined the distributed pattern of responses^16^ for each category and in each anatomical ROI (see above). To do so, we estimated response amplitudes from the GLM at each voxel for each run, which were then transformed into z-scores. We next determined all pair-wise correlations between multivoxel patterns of responses across unique combinations of runs for all categories and then averaged across run combinations. This procedure resulted in a 10x10 representational similarity matrix^17^ (RSM) for each participant. For visualization, we also computed the mean RSMs for each group (**Fig. 1D**). In the RSM, values on the diagonal show the similarity of distributed response patterns to different images of the same category while the values on the off-diagonal show the similarity of distributed response patterns for images of different categories. Examining the group RSMs reveals overall a similar pattern between signers and non-signers with a subjective difference in the category hands.

To quantify these subjective differences, we calculated the distinctiveness for each category (**Fig. 1D -bottom**). This measure captures both how similar responses are to different items of the same category and how similar (or dissimilar) they are to items of different categories^26^. Distinctiveness is higher when items within a category produce similar response patterns to one another and dissimilar response patterns to items of other categories. We first examined the distinctiveness in ventral ROIs and focused on the lateral part of VTC, because this is where face- and hand-selective regions reside^57^. We employed Bayesian statistical methods, classifying the strength of evidence according to the guidelines as in Jeffreys (1961)^58^ and in Keysers et al. (2020)^59^, where a Bayes factor (BF) between 1 and 3 is interpreted as anecdotal, BF ≥3 as moderate, BF ≥10 as strong and BF ≥30 as very strong evidence. Results reveal a highly specific modulation of the distinctiveness for hands for signers compared to non-signers. Specifically, a Bayesian independent-samples t-test provided very strong evidence that signers exhibited higher distinctiveness for hands than non-signers in the left lateral VTC (BF₁₀ = 40.2, δ = 1.02, 95% credible interval (CrI) [0.34, 1.70]; **Fig. 1E**). For all remaining categories, including faces, there was no substantial evidence for group differences (**Fig. 1E**; **Supplemental Table 1**). Interestingly, this effect in lateral VTC was specific to the left hemisphere, as there was no evidence for group differences in any category in the right lateral VTC (hands: BF₁₀ = 0.45, δ = 0.24, 95% CrI [-0.30, 0.82]; **Fig. 1E**; see **Supplementary Table 1** for full statistics). Likewise, we observed no compelling evidence for group differences in medial VTC (**Supplemental Fig. 1A,B**; **Supplemental Table 1**).

We next turned to our LOTC ROIs to examine category distinctiveness in the lateral temporal cortex. Mirroring the findings in ventral temporal cortex, we also observed evidence for higher hand distinctiveness in signers than non-signers in the left LOTC (BF₁₀ = 8.32, δ = 0.80, 95% CrI [0.18, 1.44]; **Fig. 1F**). There was little evidence for group differences in any other category, including faces (**Supplemental Table 1**). In the right LOTC, we observed only anecdotal evidence for higher hand distinctiveness in signers than non-signers (BF₁₀ = 2.89, δ = 0.64, 95% CrI [0.04, 1.27]; **Fig. 1F**).

### Signers have larger hand-selective regions than non-signers in the left hemisphere

Given these specific modulations of distributed response patterns for hands by communication with sign language, we next asked if sign language use also shapes the size of category-selective regions. Category-selective regions contain clusters of neurons that respond more to a specific category, such as faces, compared to other visual stimuli. We defined functional ROIs on participants’ individual inflated brain surfaces. Given prior behavioral evidence for differences in the perception and representation of both hands and faces in signers compared to non-signers^45,46,48,49^, we focused on face- and hand-selective regions. Category-selectivity was defined by a t-value > 3 (voxel-level) as in previous publications (e.g.^26,35,60^). The ventral hand-selective region included clusters of hand-selective voxels on the occipito-temporal sulcus (OTS) and was labeled OTS-hands^35^. The lateral hand-selective region included hand-selective clusters on the inferior temporal gyrus (ITG), middle temporal gyrus (MTG) and lateral occipitotemporal sulcus (LOS), which were summed up to the ROI LAT-hands^9^. The ventral face-selective regions were defined as face-selective clusters located on the lateral fusiform gyrus called mFus-faces and pFus-faces. The lateral face-selective regions included face-selective voxels located on the superior temporal sulcus (STS) or its sidearms^61^ and were called mSTS-faces and pSTS-faces. We first examined whether the size of hand-selective regions differed across groups. In fact, we observed that the ventral OTS-hands in the left, but not right, hemisphere was larger in signers compared to non-signers (lh: BF₁₀ = 3.61, δ = 0.68, 95% CrI [0.06, 1.33], **Fig. 2A**; rh: BF₁₀ = 0.45, δ = 0.24, 95% CrI [-0.33, 0.85], **Fig. 2A )**. Strikingly, on average, left OTS-hands was 120% larger in signers compared to non-signers. This difference between the groups was still given when the outliers (defined as values larger than 1.5 times the interquartile range) were removed from the analysis (without outliers: BF₁₀ = 4.06, δ = 0.71, 95% CrI [0.08, 1.40]).

**Figure 2.**
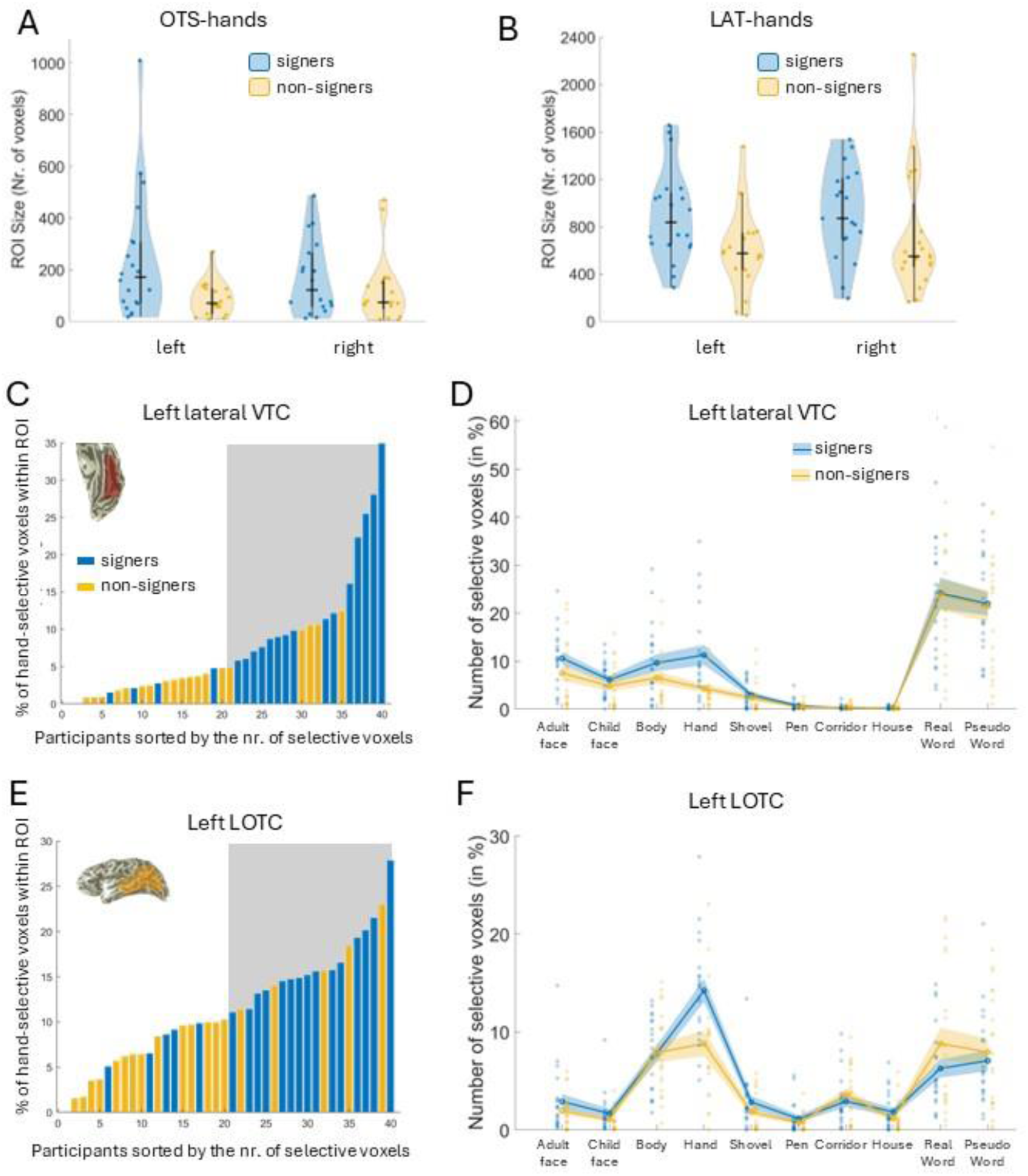
Hand-selective regio ns are larger in signers com pa red to non-signers in the left hemisphere. **A** The size of functionally defined ventral hand-selective region OTS-hand3 in the left and rght hemisphere in signers (blue) and non-signers (yellow). Colored dots = individual data points; bold horizontal black line = median; bold vertical black line = interquartile range (IQR); thin black line = 1.5 x IQR below lov«st quartile and above highest quartile. All data points above the thin black line are outliers. Numbers of includedfROIs: left OTS-hands: signers N=20. non-signers N=18; right OTS-hands: signers N=18, non-signers N=17. **B** Same as panel A but for the functionally defined lateral hand-selective region l_A T-hands. Numbers of includedfROIs; left LAT-hands: signers N=20, non-signers N=19; right LAT-hands; signers N=19, non-signers N=20. C, E Data of individual participants (20 signers. 20 non-signers) sorted from low to high by the number of hand-selective voxels in left lateral VTC (panel C) and left LOTC (panel E). The color of individual bars denotesthe group Signers are displayed in blue; non-signers are displayed in yellow. The gray background highlights the top half of the participants, showing that more signers ha>« higher numbers of hand-selective voxels in comparison to non-signers. Insets show an example of the respective Rd onthe surface, namely left lateral VTC (panel C) andleft LOTC (panel E). **D, F**Percentage of category-selective voxe Is relative to the total number of voxels in left lateral VTC (panel D) and left LOTC (panel F)l While category-selective regions displayed in A are manually defined and are restricted toclustered activation, this analysis is threshold-based, can include both clustered and unclustered cate gory-selective activations and is observer-independent. Unfilled cirdes= mean; shaded areas = standard enor of the mean; filled cirdes= individual values. Signers (N=20) are displayed in blue, non-signers(N=20)are displayed inyellow.

We next examined the size of the hand-selective region in the lateral temporal lobe (LAT-hands). Results showed that LAT-hands was larger in signers compared to non-signers in the left, but not in the right, hemisphere (lh: BF₁₀ = 3.04, δ = 0.65, 95% CrI [0.05, 1.30], **Fig. 2B**; rh: BF₁₀ = 0.58, δ = 0.32, 95% CrI [-0.24, 0.91], **Fig. 2B**). On average, left LAT-hands was 46% larger in signers compared to non-signers.

As a next step, we tested whether there were any group differences in the sizes of face-selective regions. However, consistent with the results on distinctiveness, we found no evidence for differences across groups for the sizes of face-selective regions (**Supplemental Table 2**).

Given the difference between signers and non-signers regarding the size of hand-selective regions, it is worth asking whether the amount of category-selective activation also differs for any of the other categories (besides faces). To do so, we used an observer-independent approach in which we counted the selective voxels for each of the 10 categories in our anatomically defined VTC and LOTC ROIs (**Fig. 1C**). For this approach we computed the selectivity for each voxel within an anatomical ROI for each category by contrasting the responses to a given category to that of all other categories (e.g., faces vs. all other categories). We then counted the voxels that passed our threshold for category-selectivity as in prior publications (t-value > 3, voxel level^35^) and reported the percentage of category-selective voxels relative to the overall number of voxels in the anatomical ROI.

We first tested whether this approach replicated the result of the functional region of interest analysis (**Fig. 2A,B**). Indeed, this approach revealed that signers had more hand-selective voxels compared to non-signers in left lateral VTC (BF₁₀ = 9.83, δ =0.83, 95% CrI [0.19, 1.48], **Fig. 2C,D**). Critically, this effect is also visible when examining the data on the individual level: When we ranked all 40 participants by the number of hand-selective voxels in their left lateral VTC ROI – regardless of whether they were signers or non-signers – the 20 participants with the fewest hand-selective voxels included 16 non-signers and only 4 signers (**Fig. 2C**). We next turned to the specificity of this effect and compared the number of selective voxels for all other categories across groups. No other category showed convincing evidence for group differences (**Fig. 2D**, **Supplemental Table 3**). Further, group differences for hand representations were specific to the left hemisphere, as the analysis provided no compelling evidence for group differences in the right lateral VTC (**Supplemental Fig. 2A**, **Supplemental Table 3**). Surprisingly, we found evidence for a moderate effect in left medial VTC for signers having more selective voxels for words compared to non-signers (BF₁₀ = 6.12, δ =0.76, 95% CrI [0.14, 1.43]) and anecdotal evidence for a similar effect for pseudowords (BF₁₀ = 2.12, δ =0.58, 95% CrI [0.00, 1.22], **Supplemental Fig. 2B, Supplemental Table 3**). There was no substantial evidence for group differences in the right medial VTC (**Supplemental Fig. 2C, Supplemental Table 3**).

We then turned to the LOTC ROI. Mirroring prior analyses, we found strong evidence that in the left hemisphere signers had more hand-selective voxels compared to non-signers (BF₁₀ = 10.02, δ =0.83, 95% CrI [0.19, 1.48], **Fig. 2E,F**), and no compelling evidence for group differences for any of the other categories (**Fig. 2F**). Interestingly, while non-signers had a similar distribution of hand and body-selective voxels in left LOTC (BF₁₀ =0.36, δ =0.20, 95% CrI [-0.21, 0.62]), signers showed decisive evidence for having more hand-selective voxels than body-selective voxels (BF₁₀ =614.84, δ =1.01, 95% CrI [0.52, 1.68]), indicating that sign language experience shifts the relative magnitude of different category representations in left LOTC. In the right LOTC (**Supplemental Fig. 2D**) we observed anecdotal evidence for group differences for hands (BF₁₀ =2.50, δ =0.61, 95% CrI [0.02, 1.24]), with signers having more hand-selective voxels than non-signers and for words (BF₁₀ = 2.21, δ =-0.58, 95% CrI [-1.23, 0.01]), with non-signers having more word-selective voxels than signers.

To ensure that these results do not depend on a certain threshold used to define category selectivity, we repeated our analyses with different thresholds, yielding a qualitatively similar pattern of results (**Supplemental Fig. 3**).

### Sign language experience predicts the strength of neural representations for hands in LOTC

Our prior results show that the neural representation for hands is enhanced in signers compared to non-signers. This raises the question whether experience with German Sign Language predicts the neural representations for hands within signers. We reasoned that multiple aspects of sign language experience might contribute to shaping these responses including (i) the frequency of sign language use (hours/week), (ii) the duration of sign language use (in years), and (iii) the self-assessed competence of sign language. We z-standardized each measure and averaged the three measures into a single measure of ‘sign language experience’ because we expected them to act together: For instance, someone might have used sign language for many years but with low frequency, while another person might have only recently acquired sign language but uses it with high intensity. In both cases, the overall impact on hand representations may differ from what any single factor would predict.

We first tested whether this combined measure of sign language experience can predict the number of hand-selective voxels in high-level visual cortex. We calculated a Bayesian linear regression model with sign language experience as predictor and number of hand-selective voxels as dependent variable. In fact, in left and right LOTC, the model including sign language experience was preferred over the null model (left: P(M|data) = .79, R² = .28, BF_inclusion_ = 3.69, β = 2.35, SD = 1.66, 95% CrI [−0.01, 5.10], **Fig. 3A**; right: P(M|data) = .77, R² = .27, BF_inclusion_ = 3.28, β = 1.96, SD = 1.46, 95% CrI [−0.004, 4.47], **Fig. 3A**). Thus, these models indicate that more sign language experience was associated with more hand-selective voxels in left and right LOTC. We next tested whether sign language experience was also linked to the distinctiveness for hands. This analysis revealed anecdotal evidence that sign language experience predicts distinctiveness for hands in the left hemisphere (P(M|data) = .68, R² = .22, BF_inclusion_= 2.13, β = 0.035 (SD = 0.032), 95% CrI [-0.00007, 0.09664], **Fig. 3B**) and no substantial evidence for the right hemisphere (**Supplemental Fig. 4, Supplemental Table 4**). For left and right lateral VTC, the data provided no substantial evidence for predicting hand representations from either the number of selective voxels or from the distinctiveness for hands (**Supplemental Fig. 4, Supplemental Table 4**).

**Figure 3.**
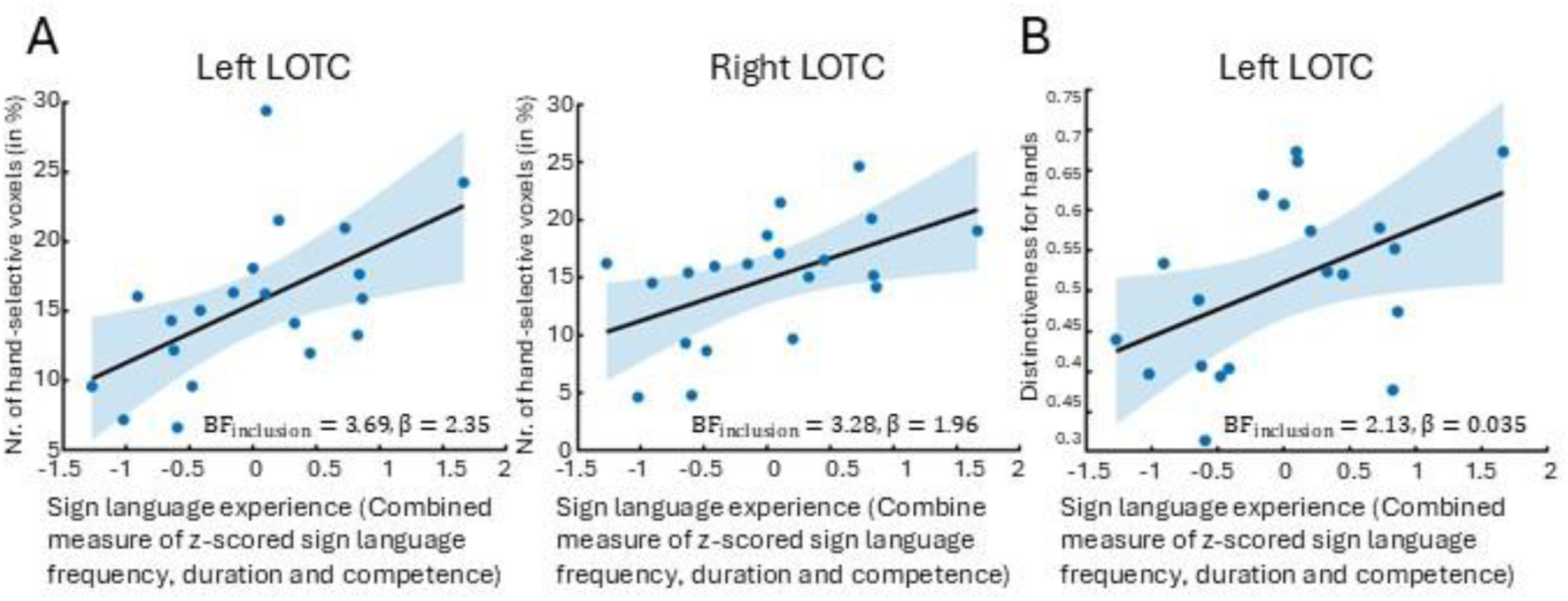
Experience with sign language predicts strength of representations for hands in LOTC. **A** Linear regression model with number of hand-selective voxels in % in left LOTC (left) and right LOTC (right) as the response variable and sign language experience as the predictor (operationalized as a combined measure of z-standardized (i)frequency of sign language use (hours/week), (ii) duration of sign language knowledge (in years) and (iii) self-assessed competence (on a scale of one (low competence) to ten (high competence)). N=20 signers. Statistics in the figure indicate the bayes factorfor inclusion of the predictor (BF_inclusion_) and the posterior mean regression coefficient (P). Full statistics can befound inthe results section. Blue dots = data points, black line= linear regression line, shaded blue area = 95% confidence band of the regression line. **B** Same as in A but predict!’ng the distinctiveness for hands in left LOTC.

### Hand representations in the ventral stream are shaped by sign language communication if acquired in adulthood

Finally, we asked if communication with German Sign Language may shape representations for hands when it was acquired in early adulthood. To address this question, we focused on the 16 participants who learned sign language in early adulthood (from now on called Late Learners). All Late Learners had acquired a sign language after they had turned 18, but they differed in the onset and duration of sign language communication (**Fig. 4A**). We first examined whether there were differences in the distinctiveness for any of the 10 categories across Late Learners and non-signers. Interestingly, our results reveal that although Late Learners acquired a sign language only in early adulthood, they showed higher distinctiveness for hands compared to non-signers in left lateral VTC (BF_10_ = 7.10, δ = 0.82, 95% CrI [0.15, 1.52], **Fig. 4B**). These effects were specific to hands as no compelling evidence for group differences in any of the other categories was found (**Fig. 4B**, **Supplemental Table 5**). In left LOTC, there was anecdotal evidence for higher distinctiveness for hands in Late Learners compared to non-signers (BF_10_ = 2.93, δ = 0.67, 95% CrI [0.03, 1.34], **Supplemental Fig. 5A**), and again no compelling evidence for group differences in any of the other categories (**Fig. 4B**, **Supplemental Fig. 5A, Supplemental Table 5**). In both cases, modulation of hand representations in Late Learners were specific to the left hemisphere as no substantial evidence was observed in right lateral VTC or right LOTC (**Supplemental Table 5**). In left medial VTC we additionally observed anecdotal evidence for higher distinctiveness for adult faces in Late Learners compared to non-signers (BF_10_ = 2.06, δ = 0.60, 95% CrI [-0.02, 1.25], **Supplemental Table 5**).

**Figure 4.**
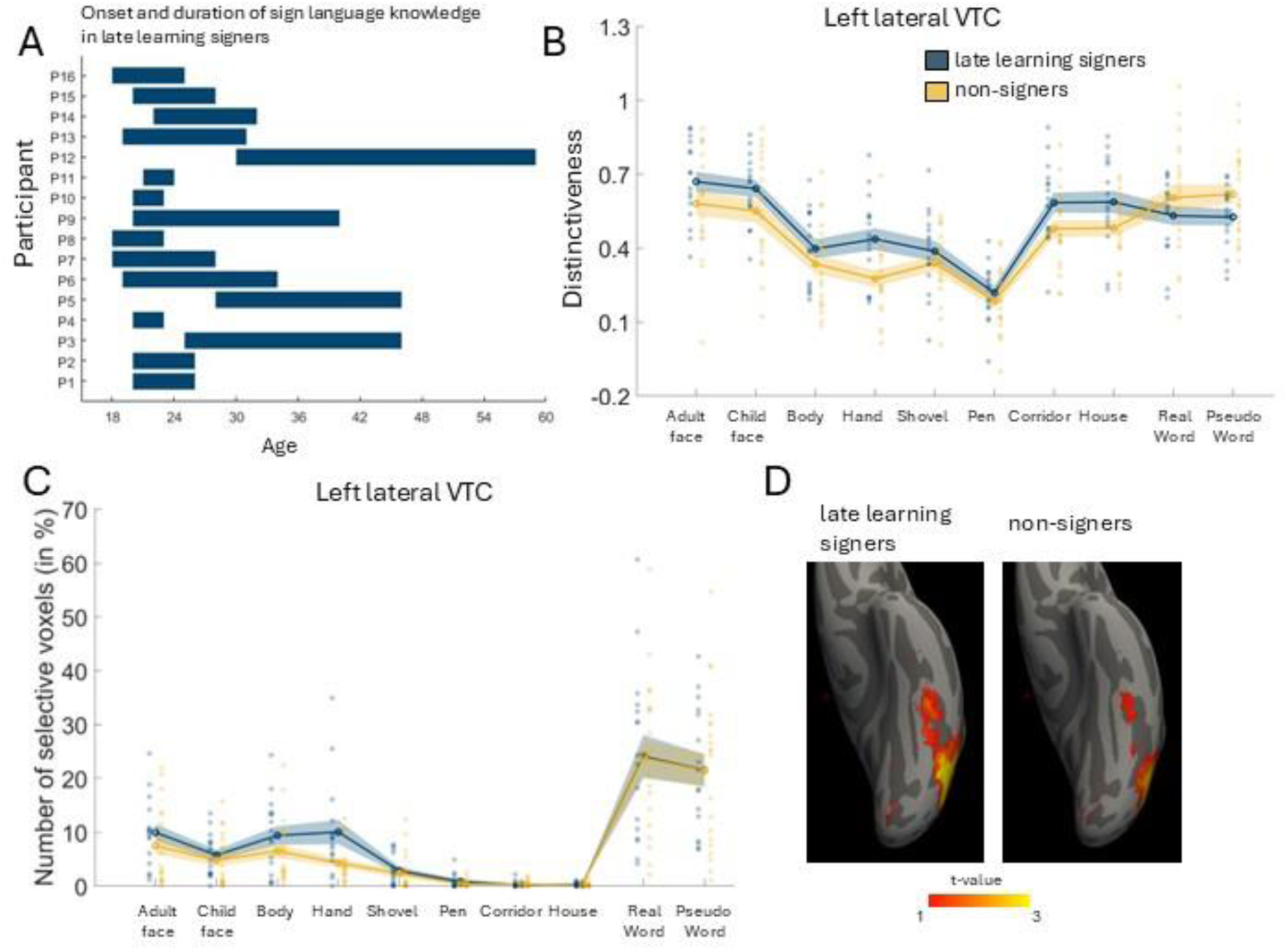
Hand representations are also shaped by sign language if acquired inearly adulthood. **A** Onsetand duration of sign language communication of late learning signers (N=16). Participant IDs are displayed on the y-axis, age on the x-axis. Bars show the onset age of learning sign language and the duration of sign language communication for each participant. **B** Distinctivenass for the ten categories in left lateral VIC in late learing signers. **C** Percentage of category-selective voxsls relative to the total number of vosels in left Lateral VTC in Late learning signers. Unfilled circles= mean; shaded areas = standard eror of themean; filled circles= individual values. **D** Mean contrast maps (limbs vs all other categories) visualized on the FreeSurfer average brain for late learning signers (left, N=16) and non-signers (right, N=20) on the ventral surface of the temporal lobe. T-values of 1 or higherare depicted (1= red and 3 orhigher = yellow). Across sl panels: dark blue: late learning signers (N=16); yellow non-signers (N=20).

Given that distributed measures are thought to be more sensitive compared to univariate analyses^26,62^, it is possible that Late Learners may show enhanced distinctiveness, but no difference in the size of category-selective regions compared to non-signers. We tested this possibility by examining the number of selective voxels for all 10 categories in our experiment. Strikingly, we found that Late Learners had a higher number of hand-selective voxels compared to the non-signers in left lateral VTC (BF_10_ = 3.92, δ = 0.72, 95% CrI [0.07, 1.40], **Fig. 4C**). To illustrate differences in hand-activation across groups on the cortical surface, we transformed individual contrast maps (contrast: hands vs all other categories) into FreeSurfer average space and visualized averaged contrast maps for both groups in left lateral VTC (**Fig. 4D**). These maps reveal more extensive activations to hands in Late Learners compared to non-signers. In left LOTC we found only anecdotal evidence for more hand-selective voxels in Late Learners compared to non-signers (BF_10_ = 2.46, δ = 0.64, 95% CrI [0.01, 1.31], **Supplemental Fig. 5B**). Critically, no effects were observed for any of the other 9 categories in left lateral VTC or left LOTC (**Fig. 4C**, **Supplemental Fig. 5B, Supplemental Table 6**). Non-signers showed anecdotal evidence for having more word-selective voxels in right LOTC, and signers showed moderate evidence for having more word-selective voxels and anecdotal evidence for having more pseudoword-selective voxels in left medial VTC (**Supplemental Table 6**). To ensure that our results do not depend on a certain threshold used to define category selectivity, we repeated our analyses with different thresholds, yielding a similar pattern of results (**Supplemental Fig. 6**).

## Discussion

Our study demonstrates that experience in communicating in German Sign Language enhances category representations in high-level visual cortex, an effect that holds in the ventral visual stream when sign language is acquired in early adulthood. Specifically, we show that (i) the distributed response patterns to hands are more distinct and hand-selective regions are larger in signers compared to non-signers, (ii) sign language experience predicts neural hand representations, and (iii) hand representations are enhanced in left lateral VTC even when sign language is acquired in early adulthood, demonstrating prolonged plasticity of high-level visual cortex.

Our results demonstrate that extensive use of German Sign Language, at approximately 20 hours per week on average, results in enhanced representations for hands in both the ventral and lateral visual streams. Critically, this effect was observed across different methodological approaches, including MVPA^16,17^, manually defined category-selective regions, and by using an observer-independent approach that identifies category-selective voxels within anatomically defined ROIs. In addition, qualitatively similar patterns of results were shown across different thresholds used to define category-selectivity. These results differ from prior expertise studies, which examine how regions selective for one category (e.g., faces) can be modified by experience with other categories (e.g., birds)^63^; our work instead focuses on effects of experience on high-level visual cortex for categories with a clustered representation.

The observed modulation of representations for hands in signers is both category-specific and more consistent in the left compared to the right hemisphere. In left lateral VTC and LOTC, our results show enhanced distributed responses to hands and a larger number of hand-selective voxels in signers compared to non-signers, but no significant effects for any of the other nine categories. Which aspect of communicating in sign language shapes the representation for hands in high-level visual cortex? We hypothesize that the increased frequency of hands in signer’s visual field is a relevant aspect leading to enhanced hand representation. Such a link between viewing behavior and neural representation is supported by developmental work showing that limb-selective regions shrink from childhood to adulthood^35,36^, a change paralleled by children looking at hands more than adults do^37,64^. As such, our results may be relevant to work in computational neuroscience testing how the visual diet shapes artificial intelligence (AI) models to achieve robust AI vision^65^. Yet, sign languages differ from spoken languages in more ways than just the frequency of hands in the visual field^42,66^. Hand and body posture, location and movement all carry semantic meaning, and these aspects may also contribute to the shown effect. Future studies can further disentangle the contribution of different aspects of sign language communication on enhanced hand representation.

Surprisingly, we did not find consistent effects of German Sign Language use on the representation of faces. Several factors may be relevant in this context. First, faces are also highly salient and extensively experienced during communication in spoken languages, so the difference between the groups might be subtle. Second, it could be that sign language does not enhance category-level face selectivity per se but rather increases the informational relevance of specific facial components and dynamics, which might have been missed by our methodological approach. Future studies employing designs that allow face representations to be targeted at a finer scale — such as fMRI adaptation^67^ — and that systematically test different aspects of face processing, may help clarify this question. Additionally it would be interesting to address these questions in a group of deaf signers, since research suggests that face processing in deaf signers might be altered in comparison to hearing signers and non-signers: Research with deaf signers has demonstrated differences in the lateralization of face processing^68,69^, which tends to be more left-lateralized compared to hearing non-signers. While some differences in lateralization compared to controls have also been observed in hearing native signers, these differed from the pattern seen in deaf signers^70^. Third, prior results suggest that face-selective regions may be less plastic in adulthood compared to childhood^24^, so it is possible that communication with sign languages may only modulate face representations when acquired during early developmental phases.

Notably, throughout our analyses we found more consistent differences in hand representations between signers and non-signers in the left hemisphere. As hand- and limb-selective regions are left-lateralized^34,35,71^ this finding shows experience-dependent effects in the dominant hemisphere. Our finding is also consistent with research showing that the left EBA supports processing of social interactions more than the right EBA^72^. Importantly, this result is also in line with the left-lateralization of motion processing in early signers compared to non-signers^73^ and of face processing in deaf signers^68,69^ and corroborates the idea that the left-lateralization of language drives the lateralization of these effects. Since the majority of signers in this study learned a sign language in early adulthood, our pattern of results is also supported by prior research suggesting that native learners of American Sign Language (ASL) show bilateral activation during sign language processing while signers who learned ASL after puberty show activation predominantly in the left hemisphere^74^. Future research examining the connectivity between hand-selective areas and language areas in signers can test how sign language experience shapes these broader networks.

Critically, our data demonstrate that experience with German Sign Language, captured by a combination of frequency, duration and self-assessed competence of sign language use, was positively linked to hand representations in LOTC. While the link between distinctiveness in left LOTC and sign language experience was only anecdotal, our results show that experience with German Sign Language was positively linked to the number of hand-selective voxels in left and right LOTC. This result raises important questions for future research. For instance, longitudinal studies in individuals acquiring a sign language could examine the timeline over which sign language experience shapes visual responses, and whether these changes emerge gradually or after a critical period of exposure. Our data also shows that sign language experience was linked to hand representations in LOTC, but not lateral VTC. We speculate that this might be related to differences in the response profile between the two regions: unlike VTC, LOTC responds stronger to dynamic than to static stimuli^75^, and is involved in processing social interactions^72^ – both properties that are relevant to sign language communication. In this context, it is noteworthy that the stimuli used in our study were static. Future studies using dynamic stimuli can test if these might evoke even larger effects.

Importantly, our results show that even in individuals who learned sign language as young adults, sign language use enhances representations for hands in left lateral VTC. For left LOTC the evidence was only anecdotal, which might be explainable by our limited sample size in the group of Late Learners (n=16). It is also possible that ventral and lateral regions differ in their timeframes of malleability. Research suggests that the ventral stream develops on a slower trajectory than the dorsal stream^76^, which brings up the question if this slower development might be associated with extended malleability in this region. Future research with larger samples that systematically vary the onset of sign language acquisition can shed light on this question. Yet, our results showing enhanced hand representations in the ventral stream in Late Learners are important for our understanding of the development and plasticity of high-level visual cortex as they show that this part of cortex exhibits a particularly prolonged plasticity. Thus, they diverge from prior findings suggesting reduced plasticity in high-level visual cortex during visual learning in adulthood^38^. Further, they call for a rethinking of high-level visual cortex development. Prior developmental work on this topic has either cross-sectionally compared children with adults^77–79^ or longitudinally followed children^20^ up to age 17^26,35^. However, the present results suggest that parts of the ventral visual stream may remain susceptible to experience-dependent change for longer than previously thought. This finding has important implications in a variety of contexts, including literacy acquisition in early adulthood^24^, and is encouraging for individuals learning sign language as adults. More broadly, our results suggest that training and learning paradigms across educational and rehabilitative settings could be designed to take advantage of this continued plasticity in the ventral visual stream.

## Methods

### Participants

We recruited 20 participants with and 20 participants without sign language knowledge. Participants from both groups had no hearing impairments, normal or corrected to normal vision and no current psychiatric or neurological diagnoses, except for one signing participant who reported mild ADHD symptoms but did not take medication. The signing participants were recruited through university sign language programs and via mailing lists for sign language interpreters and for special needs teachers. Participants without sign language knowledge were recruited through office mailing lists and newsletters and were age-matched to the group of signing participants (BF_10_ = 0.32, δ = -0,06, 95% CrI [-0.62, 0.49]). There was also no compelling evidence for group differences in sex (BF_10_= 0.94) and handedness (BF_10_= 1.52).

Non-signing participants were between 23 and 61 years old with a mean age of 35.1 (SD = 11.59). 17 participants were females, three males. 18 participants were right-handed, two left-handed. Five of the participants were students, three were research assistants and the remaining 12 participants had other professions.

Signing participants were between 22 and 62 years old with a mean age of 34.8 (SD= 13.06). 19 participants were females, one male. All participants were right-handed. 9 participants worked as sign language interpreters, one as a special-needs teacher, five were students in sign language related programs and five had other professions.

The signing participants differed in their levels of sign language knowledge and in the amount of sign language usage per week. To get an overview of these differences, participants completed a questionnaire covering information on sign language acquisition and use. Learning age of sign language varied within the group: three participants grew up with sign language since birth, one participant learned sign language as a teen and 16 participants learned sign language between the ages of 18 and 30 years. The mean learning age for all participants was 17.7 years (SD=8.33), and 21.13 years (SD=3.54) for participants who had learned sign language as an adult (Late Learners). The subsample of Late Learners also did not differ from non-signers with regard to age (BF_10_ = 0.37, δ = -0.14, 95% CrI [-0.74, 0.42]), sex (BF_10_= 0.80), handedness (BF_10_= 1.30) or education level (BF_10_= 0.43, δ = 0.08, 95% CrI [-0.64, 0.86]).

Across the whole sample of signers self-assessed frequency of sign language use ranged from 0.25 to 49 hours per week with a mean of 20.06 hours per week (SD=14.65) (**Fig. 1B**). Self-assessed competence of sign language on a scale of 1 to 10 (with 1 being the lowest competence and 10 the highest), ranged from 4 to 10 with a mean of 7,5 (SD=1.94). All signing participants used German Sign Language. Three participants stated that they also learned other sign languages, namely American Sign Language, International Sign Language and French Sign Language but that they mostly used German Sign Language.

Data from all 40 participants was included into the analyses. For four participants (2 signers, 2 non-signers) one run of the functional localizer had to be excluded due to too much motion (see below, quality criteria). For those participants the analyses were performed with two instead of three runs.

While we did not formally assess the socioeconomic status of our participants, we asked for their level of education. The distribution of the level of education was the same across groups. Two participants per group had a secondary school certificate („Realschulabschluss“ in German) and 18 participants per group had a higher educational level.

Information on participants’ race, ancestry, and ethnicity was not collected. This information is not routinely collected in many European research contexts and was not considered necessary for answering our research question. No analyses based on race, ancestry, or ethnicity were planned a priori, and demographic data collection was limited to variables directly relevant to the research question.

This study was approved by the ethical review committee of the university hospital RWTH Aachen (number: EK 22-388). Written informed consent was obtained from all participants. Participants received 60 € for participation and a reimbursement for travel costs.

### A priori power analysis

An a priori power analysis was conducted using G-Power^80^. As there were no comparable studies regarding the methods and the sample descriptions (hearing signers) during the phase of study design, the power analysis was based on^73^ and an analysis focusing on difference between motion processing between hearing signers and hearing non-signers. Bavelier et al. (2001) found greater activation in left hemisphere motion processing areas in hearing signers compared to non-signers (*F*(1,14) = 5.4, *p* = .036, *f* = 0.62). A power analysis for this effect size (α = .05, power = .80) indicated a minimum sample size of 12 per group. We recruited 20 per group to accommodate smaller potential effects and anticipated data loss due to motion artefacts.

### Procedure

Data acquisition took place at the Research Centre Juelich and included MRI scanning as well as behavioral assessments. The presented data are part of a larger assessment including additional paradigms not reported in this study. Overall, testing lasted for about 2.5 hours. For 4 participants testing was split into two sessions due to either technical errors or organizational procedures.

### Magnetic resonance imaging

### Structural MRI

We used a Siemens 3-Tesla MRI at the Research Centre Juelich with a 64-channel head coil. We implemented a T1-weighted structural sequence of approximately five minutes (resolution time (TR) = 2300ms, time to echo (TE) = 2.32ms, flip angle = 8°, resolution= 0.9 x 0.9 x 0.9 mm³) and an additional short structural sequence of approximately three minutes (TR = 2500ms, TE = 1.88ms, flip angle = 7°, resolution = 2.5 x 2.5 x 2.5 mm³), which was used to align the functional to the structural data. During structural scanning participants watched videos on YouTube or closed their eyes.

#### Functional MRI

Data was acquired with the same scanner and same-sized head coil. We recorded three functional runs of approximately five minutes each (TR = 1000ms, TE = 28ms, flip angle = 60°, resolution = 2.5 x 2.5 x 2.5 mm³).

#### Functional paradigm

During functional scanning participants took part in three runs of a category experiment modified from^56^. In this experiment, participants watched greyscale images of 10 categories (adult faces, child faces, hands, bodies without heads, shovels, pens, houses, corridors, words and pseudowords). These ten categories form 5 broader domains (faces, bodies, tools, places and characters, respectively). The images of the categories adult faces, child faces, bodies without heads, houses and corridors were taken from^56^ and we made the following changes to the remaining categories. While in the original version of paradigm, the category ‘limbs’ included both images of hands and feet, we excluded the images of feet and added additional images of hands (some of which also show arms) and renamed the category ‘hands’. The images of words and pseudowords were adapted to German. For words we selected monomorphemic nouns with a maximum of eight letters and a lexical frequency larger than 20 per million. Pseudowords were created with the program Wuggy^81^ and were matched to real words in word length and number of syllables. We replaced the two object categories with tools, including shovels and pens. As in the original experiment^56^ all images were placed on a phase-scrambled background which was created from randomly selected images. During scanning images of the same category were presented at 2hz in blocks of 4s. In between blocks, participants were presented with a grey screen for 4s as the baseline condition. Participants were instructed to fixate on a small red square at the center of the images and perform an oddball-task. This task required them to press a button when an image showing only the phase-scrambled background appeared. Performance was calculated as the percentage of correctly detected targets overall and separately for each category. Bayesian independent samples t-tests provided no compelling evidence for group differences in overall performance as well as for individual categories (**Supplemental Table 7**).

### Behavioral Measurement

#### Questionnaires

The participants were asked to fill out three questionnaires: the Edinburgh Handedness Inventory^82^ estimating their handedness, a questionnaire assessing demographic information (age, sex, profession), and a questionnaire on sign language acquisition and usage. The latter questionnaire was only filled out by signers and assessed (i) the age of sign language acquisition, (ii) the reason and approach for learning sign language, (iii) the average sign language usage per week, and (iv) the self-assessed sign language competency on a scale between 1 (not good) and 10 (very good).

### Anatomical Data Analysis

We used FreeSurfer (version 7.3.2) to segment high-resolution anatomical scans into gray and white matter and used this segmentation to generate individual inflated cortical surfaces in mrVista (https://github.com/vistalab/vistasoft/wiki/mrVista).

### fMRI Data Analysis

To analyze the functional data, we used MATLAB versions 2019a^83^ and 2017a^84^ and mrVista software package (https://github.com/vistalab/vistasoft/wiki/mrVista). FMRI-data was motion corrected for within-run motion and between-run motion. We aligned the functional data via the inplane sequence to the individual anatomical T1-weighted scan. Time courses were converted to percentage signal change by dividing each timepoint of each voxel’s data by the average response across the entire run. To quantify the contribution of each of the ten conditions

(corresponding to the ten image categories), a general linear model (GLM) was calculated separately for each voxel by convolving the stimulus presentation design with the hemodynamic response function (as implemented in SPM, www.fil.ion.ucl.ac.uk/spm).

### Quality assurance for functional data

We excluded runs with a within-run motion of > 3 functional voxels and all runs following a between-run motion value > 3.5 functional voxels. These criteria lead to the exclusion of four runs in total. After exclusion there was no significant difference in motion between the groups (t(38)=0.07, p=0.95).

### Definition of anatomical ROIs

We defined three anatomical ROIs per hemisphere on the individual brain surface of each participant: the ventral temporal cortex (VTC), separated into a medial (medial VTC) and a lateral part (lateral VTC) and the lateral occipitotemporal cortex (LOTC; including the Superior temporal Sulcus, STS). VTC ROIs (**Fig. 1C -top**) were restricted medially at the medial border of the Collateral Sulcus (CoS) and laterally at the lateral border of the OTS. The posterior border was set at the fundus of the posterior transverse CoS (ptCoS), and the anterior border was placed at the anterior tip of the mid-fusiform sulcus (MfS). We then separated the VTC into a medial and a lateral part along the MfS as in prior publications^26,35^, because we know that face- and hand-selective regions fall into the lateral part of VTC^57^, while place-selective regions fall into the medial part of VTC^57^.

For the LOTC (**Fig. 1C -bottom**), the inferior boundary of the ROI was placed along the inferior temporal gyrus (ITG), the posterior border was set posterior to the lateral occipitotemporal sulcus (LOS), and the superior border was placed along the superior outline of the STS. The anterior border was placed near the anterior tip of the MfS^3,85^. While the LOTC ROI in previous publications did not include the STS^3,85^, we included this structure because lateral face-selective regions are located in the STS^5,6,61^. We ensured that there was no overlap between anatomical ROIs.

As anatomical ROIs are based on individual anatomical landmarks, it is possible that ROI sizes differ across groups. There were no differences in ROI size (see **Supplemental Table 8**), except for the lateral VTC ROI in the left hemisphere, which was larger in non-signers compared to signers (BF₁₀ = 8.12, δ = -0.79, 95% CrI [-1.46-0.17]). Although anatomical ROI size differed between groups for this ROI, this is unlikely to confound our results as any bias introduced by this ROI size would work against our observed effects. Nevertheless, we normalized the number of category-selective voxels by the total number of voxels in each ROI to control for this difference (see below, Definition of category-selectivity in the ROIs). We also repeated the analysis in left lateral VTC with the absolute count of voxels which yielded a similar pattern of results showing more hand-selective voxels in signers compared to non-signers (category hands: BF₁₀ = 3.61, δ = 0.67, 95% CrI [0.07, 1.32]).

### Multivariate Pattern Analysis

For each participant we determined multivoxel patterns (MVP) of response amplitudes^16^ per category for the different anatomical ROIs (see above). To do so, we estimated response amplitudes from the GLM at each voxel for each run, which were then transformed into z-scores. Next, we determined all pair-wise correlations between MVPs of different run combinations (run1-run2, run2-run3, run1-run3) for all categories and then averaged across run combinations. This procedure resulted in a 10x10 RSM^17^ for each participant. We also calculated the mean RSMs across participants within each group.

### Category Distinctiveness

To quantify group differences in distributed patterns across high-level visual cortex, we determined the distinctiveness^26^ for each of the 10 categories for all participants. The distinctiveness of a given category is defined by the within-category similarity minus the average of the between category similarities from all other categories (see **Fig. 1D**-box) and can range between -2 and 2. The distinctiveness is higher when items within a category produce similar response patterns to one another and dissimilar response patterns to items of other categories.

### Definition of category-selectivity within anatomical ROIs

To compare category-selectivity between the groups, we calculated the number of selective voxels to each of the 10 categories within the anatomical ROIs (LOTC, lateral and medial VTC)^35^. To do so, each category was contrasted to all other categories including the second category from the same domain (for instance, hands vs all other categories including bodies). To define selectivity, we set a threshold at t > 3 (voxel level) as in previous publications (e.g.^35,60,86^. We then calculated the percentage of the number of selective voxels per category in relation to the total number of voxels in the respective ROI to account for possible differences in overall anatomical ROI sizes and compared these percentages between the two groups per ROI. To ensure that these results do not depend on a certain threshold used to define category selectivity, we repeated our analyses with the thresholds t > 2 and t> 4 (supplemental figures 3 and 6).

### Definition of functional ROIs (hand-selective and face-selective ROIs)

To examine the size of hand- and face-selective regions, we defined functional ROIs selective for hands and faces on participants’ individual inflated brain surfaces. ROIs were manually delineated by L.K. and controlled by M.N. For the definition we used both functional information (defined by a threshold for selectivity, see above) and individual anatomical information as in prior publications^35,60,86^. To define hand-selective regions we used the contrast ‘hands vs all other categories (including bodies)’ and for face-selective regions we contrasted the responses to adult and child faces against all other categories, because we expected no differences with regard to the responses of the two face categories. Category-selectivity was defined by a t-value > 3 (voxel level). We defined two hand-selective ROIs per hemisphere: OTS-hands and LAT-hands. Ventral clusters of hand-selective activation located on the OTS were summed up to the ROI OTS-hands. This ROI is also known as the fusiform body area^8^. Lateral clusters of hand-selective activation located on the ITG, middle temporal gyrus (MTG) and LOS, which are typically arranged in a crescent-shape^9^ were summed up to the ROI LAT-hands. This ROI is also known as the extrastriate body area^7^.

We defined four face-selective ROIs per hemisphere: mFus-faces, pFus-faces, mSTS-faces and pSTS-faces. mFus-faces and pFus-faces are face-selective clusters located on the lateral fusiform gyrus bordering the MfS^2^. Clusters aligning with the anterior tip of the MfS were defined as mFus-faces and more posterior clusters were defined as pFus-faces^2^. In combination, these ROIs are also known as the fusiform face area (FFA)^4^. mSTS-faces and pSTS-faces were located on the STS or side arms of the STS^61^. More anterior clusters, which roughly aligned with mFus-faces on the anterior to posterior axis, were defined as mSTS-faces. More posterior clusters, which roughly aligned with pFus-faces on the anterior to posterior axis, were typically located above LAT-hands and were defined as pSTS-faces^61^. If a cluster was located on more than one of the named brain structures, the assignment to one ROI was decided depending on which brain structure the major part of the cluster was located on.

### Statistics

Linear regression models and data extraction for all other analyses were performed with MATLAB version 2019a^83^. Bayesian independent samples t-tests and the Bayesian paired sample t-test were calculated in JASP version 0.98.1^87^ with the default cauchy prior of r = 0.707^88^. For comparisons of frequencies, such as the comparison of the distribution of handedness and sex, we used Bayesian contingency tables tests as implemented in JASP. In addition, some analyses were repeated with the BayesFactor package in R^89^ using the same settings, as this package readily produces formatted results tables (reported in the supplements). Next to BF_10_, which is the bayes factor in favor of the alternative hypothesis, we reported the posterior median of the standard effect size (δ) and its 95% credible interval. Positive δ values indicate higher values for signers compared to non-signers. We followed the guidelines as reported in Jeffreys (1961)^58^ and Keysers and colleagues (2020)^59^ and interpreted BF < 3 as anecdotal, BF ≥3 as moderate, BF ≥10 as strong and BF ≥30 as very strong evidence.

To predict category representations from sign language experience, we calculated bayesian linear regression models with distinctiveness or Nr. of selective voxels as response variable and sign language experience as predictor. We reasoned that multiple aspects of sign language experience might contribute to shaping responses to hands including (i) the frequency of sign language use (hours/week), (ii) the duration of sign language use (in years), and (iii) the (self-assessed) competence of sign language. We z-standardized each measure and combined the three measures into a single measure of ‘sign language experience’ because we expected them to act together. For the bayesian linear regression models we reported the posterior probability of the model (P(M|data)), the proportion of explained variance (R²), the BF for inclusion of the predictor (BF_inclusion_) and the posterior mean regression coefficient (β) with its standard deviation (SD) and 95% credible interval (95% CrI).

### Data Availability

Aggregated data and custom code to reproduce the main figures will be made available on a repository on GitHub upon publication. Raw data cannot be shared publicly due to regulations in the IRB approval.

## Acknowledgements

We thank Selina Cohnen, Theresa Heinen, Marcos Caetano, Johanna Augustin, Eva Broszy, Lilly Baksa and members of the Imaging Core Facility (ICF) at Forschungszentrum Jülich for help with data collection and we thank Aleksandra Boiko for help with data analysis. We thank SignGes for helpful discussions during planning of the project.

## Author Contributions

L.K. carried out and coordinated the project, collected and analyzed the data and wrote the manuscript. K.G. helped with project planning and corrected and reviewed the manuscript. M.S. helped with project planning and corrected and reviewed the manuscript. K.W. corrected and reviewed the manuscript. K.K. was involved in funding and corrected and reviewed the manuscript. M.N. provided funding and designed the project, contributed to the analysis pipeline and corrected and reviewed the manuscript.

