## Supplemental figures and tables for "Sign language communication enhances representations for hands in high-level visual cortex"

### Supplemental Material

#### Supplemental figures

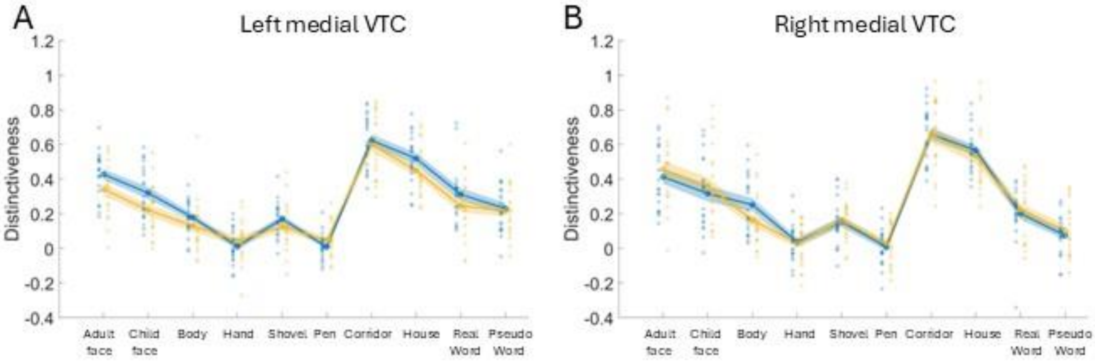


**Supplemental Figure 1. Distinctiveness in left and right medial VTC**
**A** Distinctiveness for the ten categories in the left medial VTC for signers (blue, N=20) and non-signers (yellow, N=20). Unfilled circles= mean; shaded areas = standard error of the mean; filled circles= individual distinctiveness values. **B** The same as panel A but for the right medial VTC.

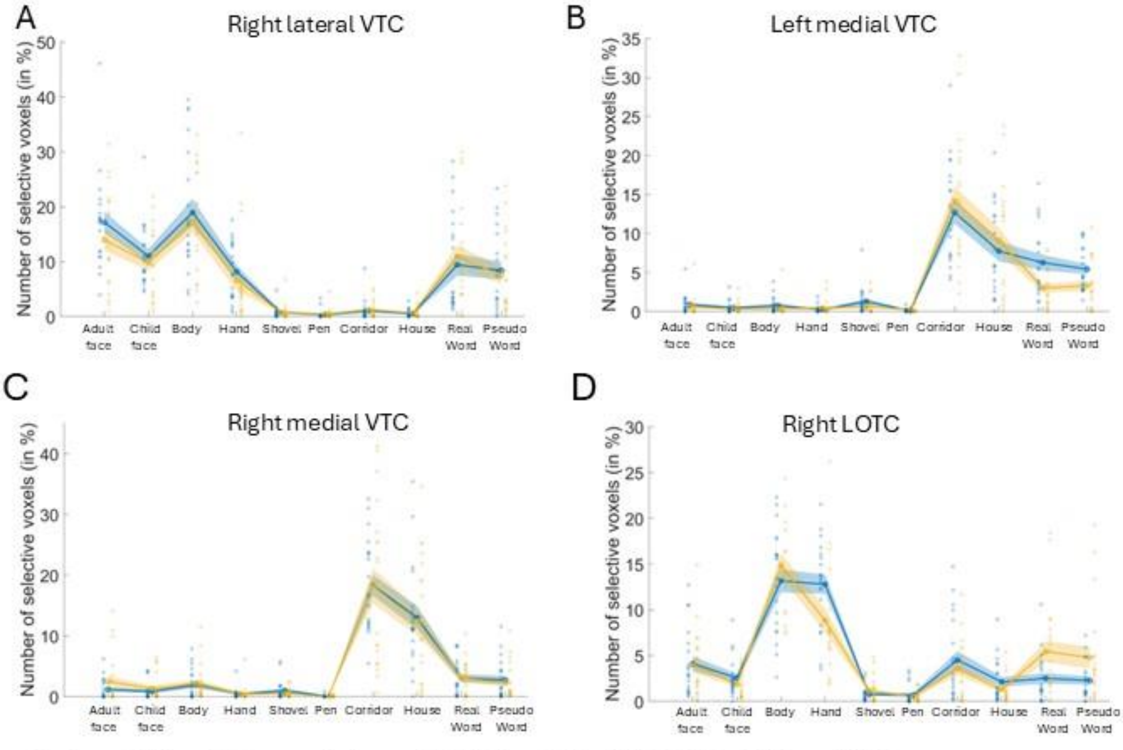


**Supplemental Figure 2. Category-selective voxels in right lateral VTC, right LOTC, left and right medial VTC**
**A** Percentage of category-selective voxels relative to the total number of voxels in right lateral VTC. Unfilled circles= mean; shaded areas = standard error of the mean; filled circles= individual values. Signers are displayed in blue; non-signers are displayed in yellow. Group sizes: signers N=20, non-signers N=20. **B** The same as panel A but for left medial VTC. **C** The same as panel A but for right medial VTC. **D** The same as panel A but for right LOTC.

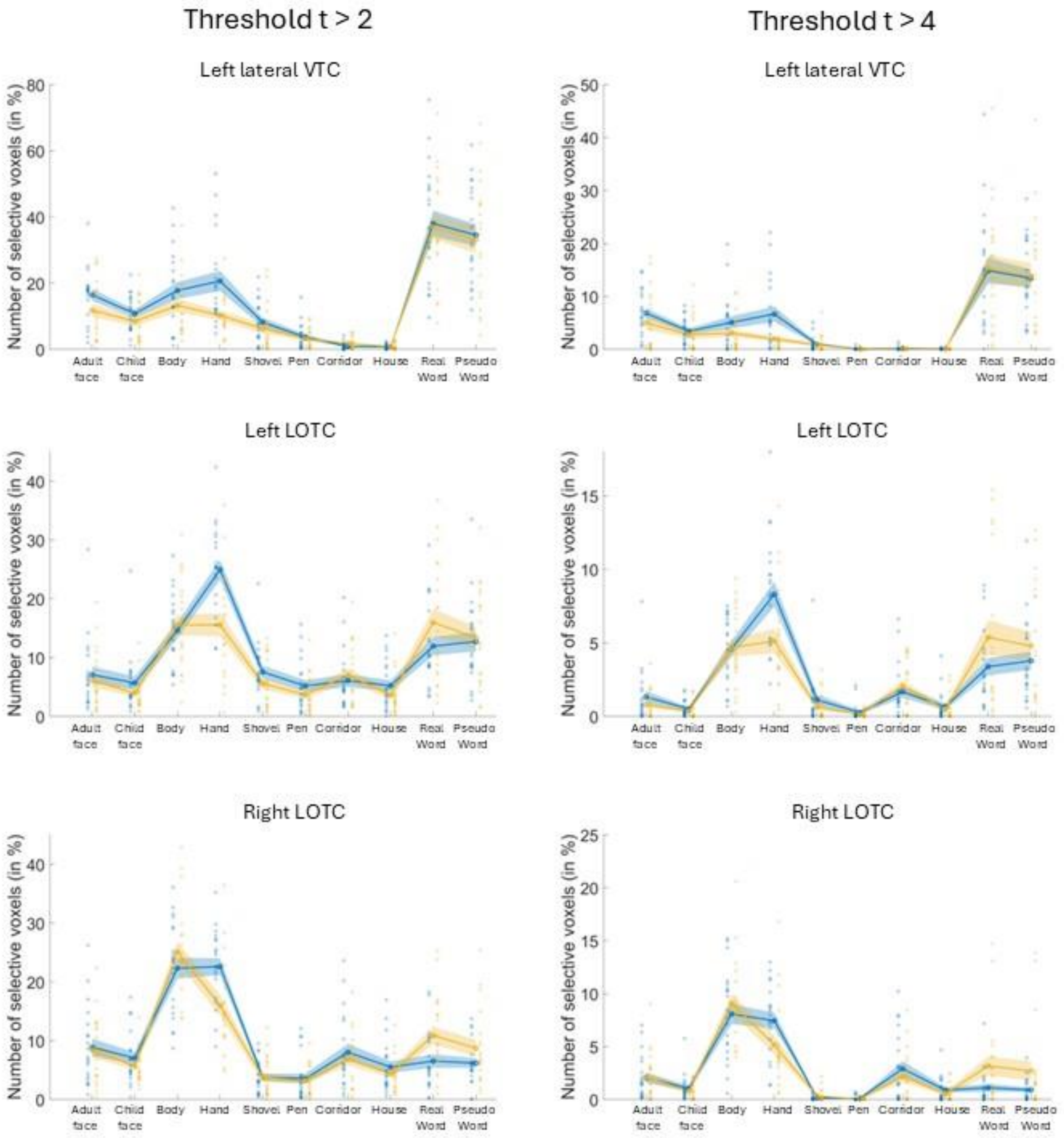


**Supplemental figure 3. Nr of selective voxel analysis using different thresholds to determine category-selectivity**
Percentage of category-selective voxels relative to the total number of voxels in left lateral VTC, left LOTC and right LOTC using different thresholds to define category-selectivity (voxel-level). Threshold $t > 2$ on the left and threshold $t > 4$ on the right. Unfilled circles= mean, shaded areas = standard error of the mean, filled circles= individual values. Signers are displayed in blue, non-signers are displayed in yellow. Group sizes: signers N=20, non-signers N=20. Statistic for the plots above: threshold > 2: left lateral VTC: hands: $BF_{10} = 10.172$, $\delta = 0.83$, 95% CrI [0.20, 1.50]; left LOTC: hands: $BF_{10} = 32.85$, $\delta = 0.99$, 95% CrI [0.33, 1.68]; right LOTC: hands: $BF_{10} = 7.67$, $\delta = 0.79$, 95% CrI [0.16, 1.45]; threshold > 4: left lateral VTC: hands: $BF_{10} = 11.00$, $\delta = 0.84$, 95% CrI [0.20, 1.51]; left LOTC: hands: $BF_{10} = 4.47$, $\delta = 0.70$, 95% CrI [0.10, 1.36]; , right LOTC: hands: $BF_{10} = 1.38$, $\delta = 0.50$, 95% CrI [-0.07, 1.13].

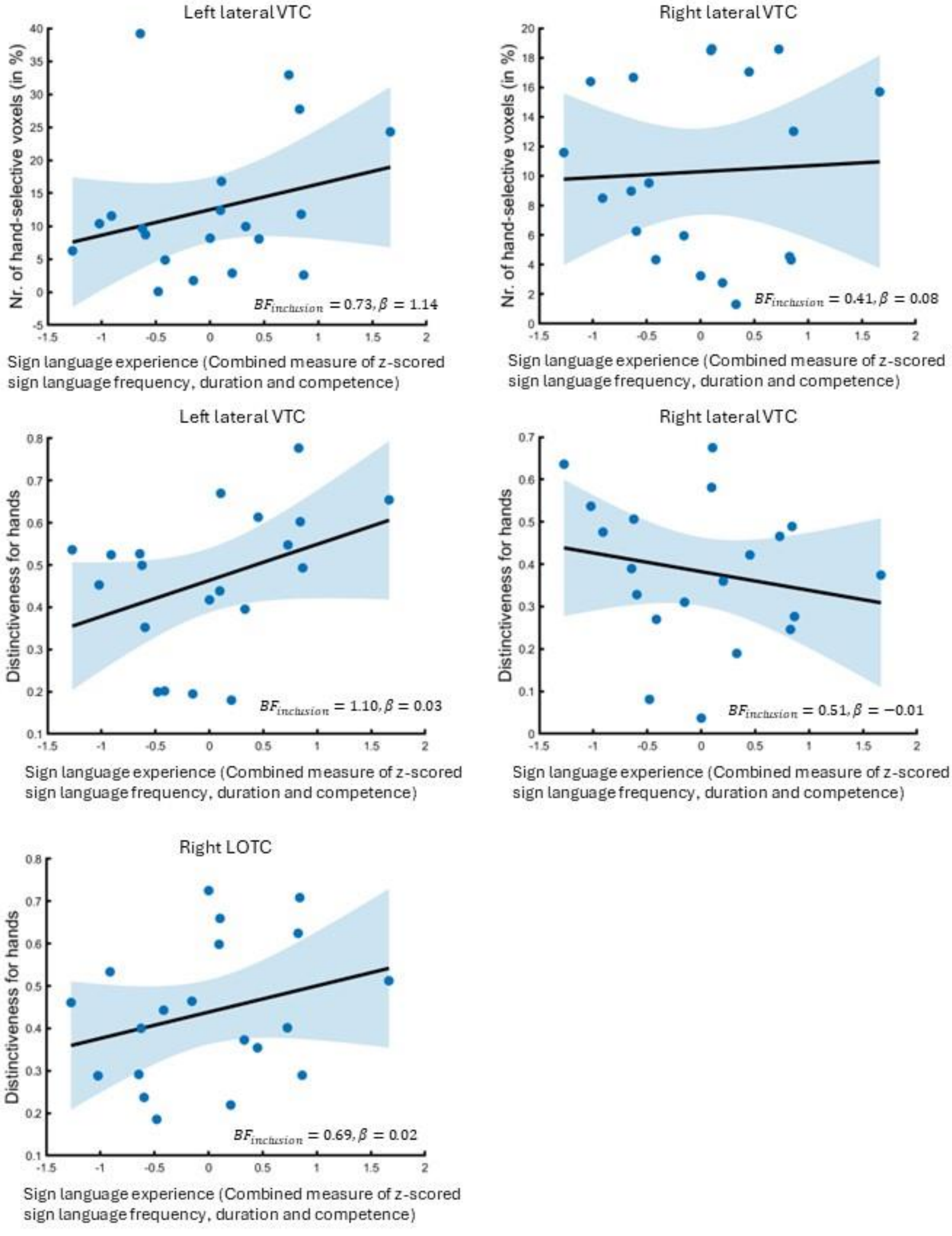


**Supplemental figure 4. Bayesian linear regression models**

Bayesian linear regression model with either the number of hand-selective voxels in % or the distinctiveness for hands as the response variable and sign language experience as the predictor (operationalized as a combined measure of z-standardized (i) frequency of sing language use (hours/week), (ii) duration of sign language knowledge (in years) and (iii) self-assessed competence (on a scale of one (low competence) to ten (high competence)). N=all 20 signers. Statistics in the figure indicate the bayes factor for inclusion of the predictor ($BF_{inclusion}$) and the posterior mean regression coefficient ($\beta$). Full statistics can be found in Supplemental Tabel 4. Blue dots = data points, black line = linear regression line, shaded blue area = 95% confidence band of the regression line.

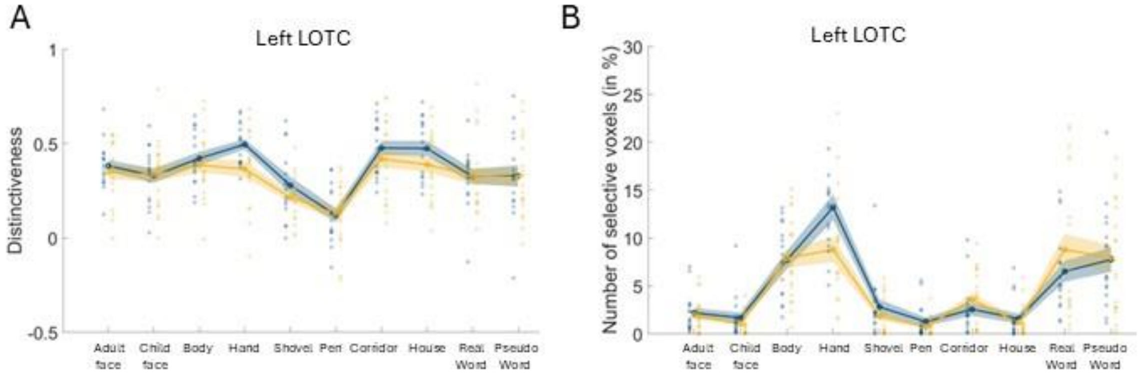


**Supplemental Figure 5. Category distinctiveness in late learning signers**
Distinctiveness (panel A) and percentage of nr. of selective voxel analysis (panel B) for the ten categories in left LOTC for late learning signers (blue, N=16) and non-signers (yellow, N=20). Unfilled circles= mean; shaded areas = standard error of the mean; filled circles= individual distinctiveness values.

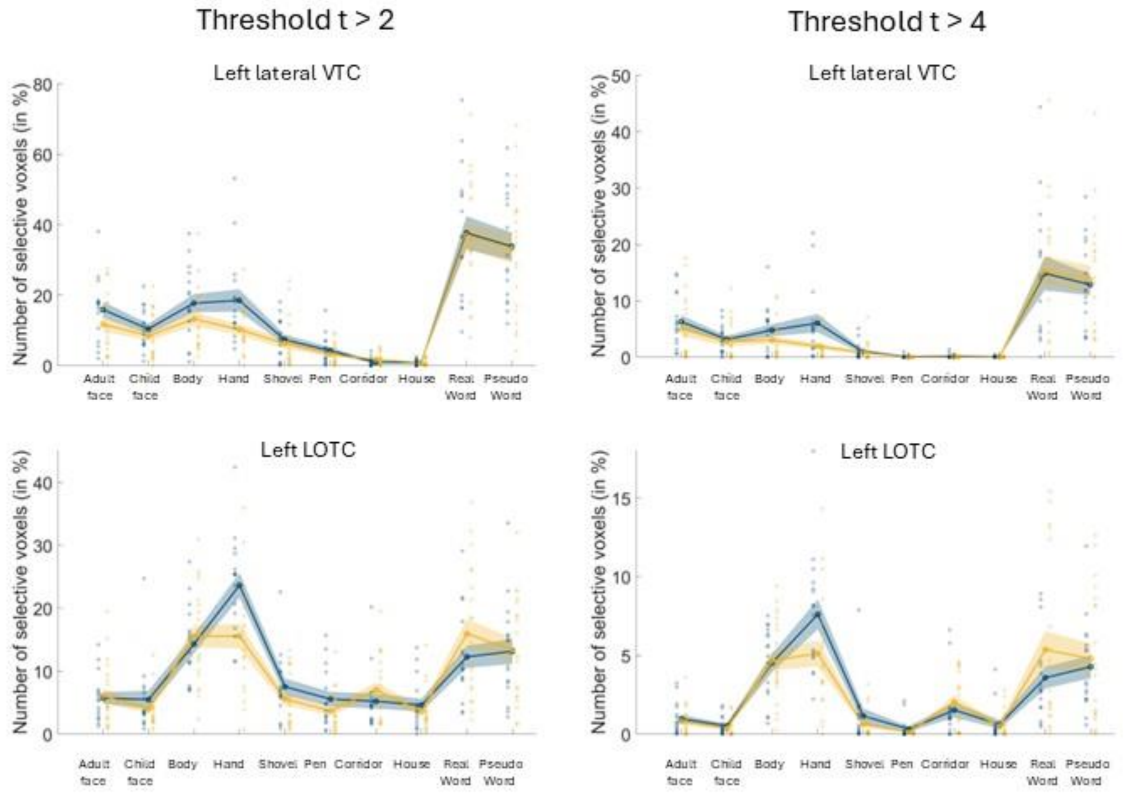


**Supplemental figure 6. Nr of selective voxel analysis with different thresholds to determine category-selectivity in late learning signers**
Percentage of category-selective voxels relative to the total number of voxels in left lateral VTC and left LOTC using different thresholds to define category-selectivity (voxel-level). Threshold t > 2 on the left and threshold t > 4 on the right. Unfilled circles = mean, shaded areas = standard error of the mean, filled circles= individual values. Signers are displayed in blue, non-signers are displayed in yellow. Group sizes: late learning signers N=16, non-signers N=20. Statistic for the plots above: threshold > 2: left lateral VTC: hands: $BF_{10}$ = 3.42, δ = 0.70, 95% CrI [0.06, 1.39]; left LOTC: hands: $BF_{10}$ = 6.46, δ = 0.80, 95% CrI [0.14, 1.51]; threshold > 4: left lateral VTC: hands: $BF_{10}$ = 4.15, δ = 0.73, 95% CrI [0.08, 1.42]; left LOTC: hands: $BF_{10}$ = 1.38, δ = 0.52, 95% CrI [-0.08, 1.19].

#### Supplemental tables

*Left LOTC:*

| Category | Signers M (SD) | Non-signers M (SD) | $BF_{10}$ | δ [95% CrI] |
|---|---|---|---|---|
| Adult Faces | 0.39 (0.13) | 0.35 (0.16) | 0.40 | 0.20 [-0.33, 0.77] |
| Child Faces | 0.33 (0.14) | 0.34 (0.18) | 0.31 | -0.01 [-0.56, 0.52] |
| Bodies | 0.42 (0.15) | 0.39 (0.17) | 0.36 | 0.15 [-0.40, 0.71] |
| Hands | 0.51 (0.11) | 0.37 (0.19) | 8.32 | 0.80 [0.18, 1.44] |
| Shovels | 0.28 (0.17) | 0.22 (0.12) | 0.75 | 0.38 [-0.17, 0.99] |
| Pens | 0.14 (0.14) | 0.13 (0.16) | 0.31 | 0.05 [-0.51, 0.61] |
| Corridors | 0.47 (0.16) | 0.42 (0.20) | 0.41 | 0.20 [-0.35, 0.77] |
| Houses | 0.46 (0.15) | 0.39 (0.18) | 0.66 | 0.34 [-0.20, 0.95] |
| Words | 0.32 (0.14) | 0.33 (0.20) | 0.31 | -0.04 [-0.61, 0.51] |
| Pseudowords | 0.31 (0.21) | 0.32 (0.21) | 0.31 | -0.04 [-0.59, 0.52] |

*Right LOTC:*

| Category | Signers M (SD) | Non-signers M (SD) | $BF_{10}$ | δ [95% CrI] |
|---|---|---|---|---|
| Adult Faces | 0.40 (0.16) | 0.40 (0.18) | 0.31 | 0.00 [-0.53, 0.55] |
| Child Faces | 0.36 (0.15) | 0.38 (0.18) | 0.33 | -0.09 [-0.63, 0.45] |
| Bodies | 0.58 (0.18) | 0.50 (0.13) | 0.86 | 0.40 [-0.14, 1.03] |
| Hands | 0.44 (0.16) | 0.31 (0.17) | 2.89 | 0.64 [0.04, 1.27] |
| Shovels | 0.20 (0.12) | 0.21 (0.10) | 0.34 | -0.13 [-0.70, 0.41] |
| Pens | 0.15 (0.13) | 0.17 (0.19) | 0.33 | -0.09 [-0.66, 0.47] |
| Corridors | 0.54 (0.17) | 0.46 (0.20) | 0.65 | 0.35 [-0.20, 0.94] |
| Houses | 0.52 (0.14) | 0.45 (0.12) | 0.76 | 0.38 [-0.17, 0.98] |
| Words | 0.18 (0.14) | 0.20 (0.16) | 0.34 | -0.12 [-0.69, 0.43] |
| Pseudowords | 0.11 (0.20) | 0.15 (0.15) | 0.39 | -0.19 [-0.77, 0.36] |

*Left lateral VTC:*

| Category | Signers M (SD) | Non-signers M (SD) | $BF_{10}$ | δ [95% CrI] |
|---|---|---|---|---|
| Adult Faces | 0.68 (0.16) | 0.58 (0.22) | 0.86 | 0.41 [-0.15, 1.00] |
| Child Faces | 0.66 (0.13) | 0.55 (0.21) | 1.35 | 0.50 [-0.07, 1.12] |
| Bodies | 0.40 (0.15) | 0.34 (0.18) | 0.57 | 0.31 [-0.23, 0.91] |
| Hands | 0.46 (0.17) | 0.28 (0.15) | 40.30 | 1.02 [0.34, 1.70] |
| Shovels | 0.39 (0.15) | 0.34 (0.14) | 0.54 | 0.29 [-0.26, 0.88] |
| Pens | 0.23 (0.12) | 0.18 (0.13) | 0.52 | 0.28 [-0.27, 0.87] |
| Corridors | 0.57 (0.15) | 0.48 (0.16) | 1.38 | 0.51 [-0.06, 1.13] |
| Houses | 0.56 (0.18) | 0.48 (0.14) | 0.89 | 0.42 [-0.14, 1.02] |
| Words | 0.55 (0.16) | 0.60 (0.24) | 0.41 | -0.20 [-0.78, 0.35] |
| Pseudowords | 0.55 (0.15) | 0.62 (0.18) | 0.62 | -0.33 [-0.93, 0.22] |

*Right lateral VTC:*

| Category | Signers M (SD) | Non-signers M (SD) | $BF_{10}$ | δ [95% CrI] |
|---|---|---|---|---|
| Adult Faces | 0.80 (0.12) | 0.72 (0.22) | 0.73 | 0.37 [-0.18, 0.95] |
| Child Faces | 0.77 (0.13) | 0.68 (0.19) | 0.83 | 0.40 [-0.15, 1.00] |
| Bodies | 0.49 (0.17) | 0.48 (0.19) | 0.31 | 0.05 [-0.48, 0.62] |
| Hands | 0.38 (0.17) | 0.33 (0.18) | 0.45 | 0.24 [-0.30, 0.82] |
| Shovels | 0.31 (0.15) | 0.34 (0.10) | 0.35 | -0.14 [-0.72, 0.39] |
| Pens | 0.27 (0.16) | 0.24 (0.20) | 0.36 | 0.15 [-0.40, 0.72] |
| Corridors | 0.57 (0.14) | 0.50 (0.16) | 0.79 | 0.39 [-0.15, 0.99] |
| Houses | 0.53 (0.17) | 0.51 (0.13) | 0.34 | 0.13 [-0.42, 0.68] |
| Words | 0.33 (0.17) | 0.36 (0.18) | 0.36 | -0.15 [-0.72, 0.40] |
| Pseudowords | 0.34 (0.24) | 0.35 (0.18) | 0.31 | -0.03 [-0.60, 0.52] |

*Left medial VTC:*

| Category | Signers M (SD) | Non-signers M (SD) | $BF_{10}$ | δ [95% CrI] |
|---|---|---|---|---|
| Adult Faces | 0.43 (0.13) | 0.34 (0.16) | 1.19 | 0.47 [-0.09, 1.07] |
| Child Faces | 0.32 (0.16) | 0.22 (0.15) | 1.58 | 0.53 [-0.04, 1.15] |
| Bodies | 0.18 (0.10) | 0.13 (0.16) | 0.53 | 0.29 [-0.24, 0.90] |
| Hands | 0.01 (0.10) | 0.04 (0.12) | 0.42 | -0.21 [-0.77, 0.32] |
| Shovels | 0.17 (0.10) | 0.13 (0.14) | 0.53 | 0.28 [-0.26, 0.87] |
| Pens | 0.01 (0.08) | 0.05 (0.09) | 0.67 | -0.35 [-0.96, 0.22] |
| Corridors | 0.62 (0.15) | 0.60 (0.18) | 0.33 | 0.11 [-0.43, 0.67] |
| Houses | 0.52 (0.14) | 0.45 (0.15) | 0.72 | 0.37 [-0.19, 0.97] |
| Words | 0.31 (0.16) | 0.25 (0.18) | 0.55 | 0.30 [-0.24, 0.90] |
| Pseudowords | 0.23 (0.14) | 0.22 (0.15) | 0.32 | 0.08 [-0.47, 0.64] |

*Right medial VTC:*

| Category | Signers M (SD) | Non-signers M (SD) | $BF_{10}$ | δ [95% CrI] |
|---|---|---|---|---|
| Adult Faces | 0.41 (0.18) | 0.46 (0.22) | 0.37 | -0.17 [-0.73, 0.35] |
| Child Faces | 0.32 (0.20) | 0.36 (0.22) | 0.37 | -0.17 [-0.72, 0.37] |
| Bodies | 0.25 (0.17) | 0.16 (0.16) | 0.93 | 0.42 [-0.13, 1.05] |
| Hands | 0.04 (0.11) | 0.03 (0.12) | 0.31 | 0.05 [-0.49, 0.60] |
| Shovels | 0.15 (0.13) | 0.16 (0.11) | 0.31 | -0.04 [-0.60, 0.50] |
| Pens | 0.01 (0.11) | 0.03 (0.12) | 0.35 | -0.14 [-0.71, 0.42] |
| Corridors | 0.66 (0.16) | 0.66 (0.20) | 0.31 | 0.00 [-0.54, 0.56] |
| Houses | 0.57 (0.12) | 0.53 (0.20) | 0.36 | 0.15 [-0.39, 0.71] |
| Words | 0.20 (0.16) | 0.22 (0.17) | 0.33 | -0.10 [-0.66, 0.45] |
| Pseudowords | 0.07 (0.11) | 0.10 (0.15) | 0.37 | -0.17 [-0.74, 0.38] |

**Supplemental Table 1. Bayesian independent-samples t-tests for Distinctiveness for all anatomical ROIs**

N of both groups = 20. $BF_{10}$ = bayes factor in favor of the alternative hypothesis; δ = posterior standardized effect size; CrI = credible interval. Positive δ values indicate higher values for signers compared to non-signers. Bayes factors were calculated using a cauchy prior with r = 0.707.

Hands-selective ROIs:

| Category | Hemisphere | N signers | N non-signers | Signers M (SD) | Non-signers M (SD) | $BF_{10}$ | δ [95% CrI] |
|---|---|---|---|---|---|---|---|
| OTS-Hands | Lh | 20 | 18 | 238.65 (244.97) | 86.78 (65.62) | 3.61 | 0.68 [0.06, 1.33] |
| OTS-Hands | Rh | 18 | 17 | 166.78 (143.04) | 123.59 (134.96) | 0.45 | 0.24 [-0.33, 0.85] |
| LAT-Hands | Lh | 20 | 19 | 898.50 (385.57) | 610.11 (347.31) | 3.04 | 0.65 [0.05, 1.30] |
| LAT-Hands | Rh | 19 | 20 | 916.79 (382.55) | 732.20 (516.58) | 0.58 | 0.32 [-0.24, 0.91] |

Face-selective ROIs:

| Category | Hemisphere | N signers | N non-signers | Signers M (SD) | Non-signers M (SD) | $BF_{10}$ | δ [95% CrI] |
|---|---|---|---|---|---|---|---|
| mFus-Faces | Lh | 19 | 15 | 117.63 (88.13) | 89.20 (77.83) | 0.48 | 0.25 [-0.32, 0.90] |
| mFus-Faces | Rh | 18 | 17 | 148.67 (103.19) | 190.12 (142.44) | 0.48 | -0.25 [-0.87, 0.33] |
| pFus-Faces | Lh | 16 | 17 | 171.31 (118.44) | 136.41 (91.84) | 0.47 | 0.25 [-0.34, 0.88] |
| pFus-Faces | Rh | 19 | 19 | 294.63 (173.98) | 245.95 (186.34) | 0.41 | 0.21 [-0.35, 0.80] |
| mSTS-Faces | Lh | 8 | 9 | 25.50 (25.40) | 44.33 (32.19) | 0.75 | -0.42 [-1.37, 0.33] |
| mSTS-Faces | Rh | 8 | 13 | 87.25 (124.43) | 90.54 (76.73) | 0.40 | -0.02 [-0.75, 0.70] |
| pSTS-Faces | Lh | 14 | 15 | 226.07 (267.76) | 121.93 (225.47) | 0.57 | 0.31 [-0.32, 1.00] |
| pSTS-Faces | Rh | 17 | 15 | 156.53 (132.16) | 132.13 (111.81) | 0.38 | 0.15 [-0.45, 0.77] |

**Supplemental Table 2. Bayesian independent-samples t-tests for the functional ROI Analysis for hands and faces**

$BF_{10}$ = bayes factor in favor of the alternative hypothesis; δ = posterior standardized effect size; CrI = credible interval. Positive δ values indicate higher values for signers compared to non-signers. Bayes factors were calculated using a cauchy prior with r = 0.707.

*Left LOTC:*

| Category | Signers M (SD) | Non-signers M (SD) | $BF_{10}$ | δ [95% CrI] |
|---|---|---|---|---|
| Adult Faces | 2.88 (3.43) | 1.95 (1.81) | 0.49 | 0.26 [-0.28, 0.83] |
| Child Faces | 1.68 (2.02) | 1.05 (1.10) | 0.56 | 0.30 [-0.23, 0.89] |
| Bodies | 7.80 (3.69) | 7.87 (4.29) | 0.31 | -0.01 [-0.55, 0.55] |
| Hands | 14.22 (5.38) | 8.80 (5.83) | 10.02 | 0.83 [0.19, 1.48] |
| Shovels | 2.83 (2.86) | 1.83 (1.91) | 0.60 | 0.32 [-0.24, 0.91] |
| Pens | 1.14 (1.63) | 0.74 (0.92) | 0.44 | 0.23 [-0.32, 0.81] |
| Corridors | 2.91 (2.67) | 3.58 (2.92) | 0.39 | -0.18 [-0.75, 0.36] |
| Houses | 1.84 (1.96) | 1.16 (1.62) | 0.54 | 0.30 [-0.25, 0.88] |
| Words | 6.27 (4.21) | 8.80 (7.30) | 0.63 | -0.33 [-0.93, 0.23] |
| Pseudowords | 7.06 (4.82) | 7.93 (6.01) | 0.34 | -0.12 [-0.69, 0.43] |

*Right LOTC:*

| Category | Signers M (SD) | Non-signers M (SD) | $BF_{10}$ | δ [95% CrI] |
|---|---|---|---|---|
| Adult Faces | 4.11 (3.63) | 3.96 (3.71) | 0.31 | 0.03 [-0.52, 0.56] |
| Child Faces | 2.55 (2.13) | 1.85 (1.46) | 0.56 | 0.30 [-0.23, 0.89] |
| Bodies | 13.16 (5.66) | 14.86 (5.73) | 0.44 | -0.23 [-0.81, 0.33] |
| Hands | 12.79 (4.74) | 8.82 (5.92) | 2.50 | 0.61 [0.02, 1.24] |
| Shovels | 0.78 (0.81) | 1.14 (1.43) | 0.46 | -0.25 [-0.84, 0.29] |
| Pens | 0.69 (1.13) | 0.51 (0.61) | 0.36 | 0.16 [-0.39, 0.73] |
| Corridors | 4.51 (4.44) | 3.53 (3.16) | 0.40 | 0.20 [-0.34, 0.78] |
| Houses | 2.13 (2.34) | 1.30 (1.41) | 0.64 | 0.34 [-0.21, 0.93] |
| Words | 2.51 (2.63) | 5.44 (5.15) | 2.21 | -0.58 [-1.23, 0.01] |
| Pseudowords | 2.27 (2.03) | 4.78 (5.37) | 1.36 | -0.50 [-1.13, 0.07] |

*Left lateral VTC:*

| Category | Signers M (SD) | Non-signers M (SD) | $BF_{10}$ | δ [95% CrI] |
|---|---|---|---|---|
| Adult Faces | 10.57 (5.82) | 7.50 (7.07) | 0.75 | 0.37 [-0.18, 0.96] |
| Child Faces | 6.11 (3.78) | 4.79 (4.76) | 0.45 | 0.23 [-0.29, 0.82] |
| Bodies | 9.63 (7.92) | 6.50 (5.84) | 0.68 | 0.35 [-0.19, 0.97] |
| Hands | 11.24 (9.51) | 4.29 (3.68) | 9.83 | 0.83 [0.19, 1.48] |
| Shovels | 2.94 (2.72) | 2.33 (3.55) | 0.36 | 0.14 [-0.40, 0.71] |
| Pens | 0.67 (1.24) | 0.39 (0.64) | 0.42 | 0.22 [-0.33, 0.80] |
| Corridors | 0.26 (0.53) | 0.38 (0.54) | 0.38 | -0.18 [-0.74, 0.37] |
| Houses | 0.23 (0.32) | 0.14 (0.31) | 0.44 | 0.23 [-0.31, 0.81] |
| Words | 24.14 (14.67) | 23.80 (14.30) | 0.31 | 0.02 [-0.53, 0.58] |
| Pseudowords | 21.94 (10.93) | 21.40 (14.65) | 0.31 | 0.04 [-0.52, 0.59] |

*Right lateral VTC:*

| Category | Signers M (SD) | Non-signers M (SD) | $BF_{10}$ | δ [95% CrI] |
|---|---|---|---|---|
| Adult Faces | 17.17 (8.77) | 14.00 (8.81) | 0.52 | 0.28 [-0.26, 0.85] |
| Child Faces | 11.02 (5.78) | 9.79 (6.40) | 0.36 | 0.15 [-0.37, 0.72] |
| Bodies | 18.96 (11.20) | 17.34 (9.12) | 0.34 | 0.12 [-0.41, 0.70] |
| Hands | 8.15 (5.32) | 6.60 (8.31) | 0.38 | 0.17 [-0.37, 0.74] |
| Shovels | 0.74 (1.05) | 0.79 (1.50) | 0.31 | -0.03 [-0.59, 0.51] |
| Pens | 0.34 (0.87) | 0.43 (1.03) | 0.32 | -0.07 [-0.64, 0.49] |
| Corridors | 1.05 (1.96) | 1.20 (1.55) | 0.32 | -0.06 [-0.61, 0.48] |
| Houses | 0.53 (0.75) | 0.66 (1.15) | 0.33 | -0.10 [-0.67, 0.43] |
| Words | 9.42 (8.97) | 10.92 (9.30) | 0.34 | -0.12 [-0.69, 0.42] |
| Pseudowords | 8.39 (7.08) | 7.54 (7.15) | 0.33 | 0.10 [-0.46, 0.65] |

*Left medial VTC:*

| Category | Signers M (SD) | Non-signers M (SD) | $BF_{10}$ | δ [95% CrI] |
|---|---|---|---|---|
| Adult Faces | 0.89 (1.24) | 0.62 (1.37) | 0.36 | 0.15 [-0.39, 0.70] |
| Child Faces | 0.47 (0.79) | 0.35 (0.76) | 0.34 | 0.12 [-0.40, 0.68] |
| Bodies | 0.77 (1.03) | 0.38 (1.21) | 0.50 | 0.27 [-0.26, 0.87] |
| Hands | 0.24 (0.49) | 0.45 (1.05) | 0.40 | -0.20 [-0.75, 0.34] |
| Shovels | 1.28 (1.78) | 0.63 (1.12) | 0.66 | 0.34 [-0.21, 0.94] |
| Pens | 0.10 (0.22) | 0.27 (0.62) | 0.50 | -0.28 [-0.87, 0.29] |
| Corridors | 12.71 (6.51) | 14.19 (8.23) | 0.36 | -0.15 [-0.71, 0.39] |
| Houses | 7.74 (5.69) | 9.13 (7.01) | 0.37 | -0.16 [-0.73, 0.37] |
| Words | 6.27 (4.51) | 2.99 (2.57) | 6.12 | 0.76 [0.14, 1.43] |
| Pseudowords | 5.43 (2.88) | 3.31 (3.09) | 2.12 | 0.58 [0.00, 1.22] |

*Right medial VTC:*

| Category | Signers M (SD) | Non-signers M (SD) | $BF_{10}$ | δ [95% CrI] |
|---|---|---|---|---|
| Adult Faces | 1.17 (1.73) | 2.48 (4.24) | 0.59 | -0.32 [-0.91, 0.21] |
| Child Faces | 0.85 (1.46) | 1.33 (2.20) | 0.40 | -0.20 [-0.76, 0.34] |
| Bodies | 1.96 (2.43) | 2.14 (3.02) | 0.31 | -0.05 [-0.60, 0.50] |
| Hands | 0.43 (0.93) | 0.45 (1.37) | 0.31 | -0.01 [-0.56, 0.52] |
| Shovels | 0.99 (1.68) | 0.53 (0.78) | 0.50 | 0.27 [-0.28, 0.85] |
| Pens | 0.06 (0.19) | 0.06 (0.12) | 0.31 | 0.03 [-0.52, 0.60] |
| Corridors | 18.32 (7.64) | 18.23 (13.01) | 0.31 | 0.01 [-0.54, 0.56] |
| Houses | 13.00 (8.48) | 12.40 (10.30) | 0.31 | 0.05 [-0.49, 0.59] |
| Words | 3.04 (2.93) | 3.07 (2.83) | 0.31 | -0.00 [-0.56, 0.56] |
| Pseudowords | 2.72 (3.28) | 2.31 (3.01) | 0.33 | 0.10 [-0.45, 0.66] |

**Supplemental Table 3. Bayesian independent-samples t-tests for Nr. of selective voxel analysis for all anatomical ROIs**

N of both groups = 20. $BF_{10}$ = bayes factor in favor of the alternative hypothesis; δ = posterior standardized effect size; CrI = credible interval. Positive δ values indicate higher values for signers compared to non-signers. Bayes factors were calculated using a cauchy prior with r = 0.707.

Nr. of selective Voxels

| ROI | Model | | Predictor | | | |
|---|---|---|---|---|---|---|
| | P(M\|data) | $R^2$ | $BF_{inclusion}$ | Posterior mean (β) | SD | 95% CrI |
| Left LOTC | 0.7870 | 0.2817 | 3.6941 | 2.3499 | 1.6585 | -0.0124, 5.0954 |
| Left lateral VTC | 0.4233 | 0.0901 | 0.7341 | 1.1350 | 2.0386 | -1.6581, 5.9535 |
| Right LOTC | 0.7664 | 0.2695 | 3.2806 | 1.9580 | 1.4583 | -0.0044, 4.4699 |
| Right lateral VTC | 0.2882 | 0.0031 | 0.4050 | 0.0760 | 0.7400 | -1.8916, 2.0506 |

Distinctiveness

| ROI | Model | | Predictor | | | |
|---|---|---|---|---|---|---|
| | P(M\|data) | $R^2$ | $BF_{inclusion}$ | Posterior mean (β) | SD | 95% CrI |
| Left LOTC | 0.6810 | 0.2230 | 2.1346 | 0.0350 | 0.0321 | 0.0000, 0.0957 |
| Left lateral VTC | 0.5246 | 0.1441 | 1.1034 | 0.0329 | 0.0437 | -0.0028, 0.1368 |
| Right LOTC | 0.4085 | 0.0817 | 0.6907 | 0.0179 | 0.0341 | -0.0167, 0.1177 |
| Richt lateral VTC | 0.3378 | 0.381 | 0.5101 | -0.0103 | 0.0290 | -0.1038, 0.0389 |

**Supplemental Table 4. Statistics for the bayesian linear regression models**
Statistics for the linear regression models with nr. of hand-selective voxels in % (top) or distinctiveness (bottom) as the response variable and sign language experience as the predictor (operationalized as a combined measure of z-standardized (i) frequency of sign language use (hours/week), (ii) duration of sign language knowledge (in years) and (iii) self-assessed competence (on a scale of one (low competence) to ten (high competence)). N=all 20 signers. P(M|data) = posterior probability of the model; $R^2$ = proportion of explained variance; $BF_{inclusion}$= Bayes factor for inclusion of the predictor; β = posterior mean regression coefficient; SD = standard deviation of β; 95% CrI = 95% credible interval of β.

*Left LOTC:*

| Category | Late Learners M (SD) | Non-signers M (SD) | $BF_{10}$ | δ [95% CrI] |
|---|---|---|---|---|
| Adult Faces | 0.38 (0.12) | 0.35 (0.16) | 0.38 | 0.16 [-0.40, 0.75] |
| Child Faces | 0.33 (0.15) | 0.34 (0.18) | 0.32 | -0.03 [-0.60, 0.54] |
| Bodies | 0.42 (0.14) | 0.39 (0.17) | 0.39 | 0.17 [-0.39, 0.79] |
| Hands | 0.50 (0.11) | 0.37 (0.19) | 2.93 | 0.67 [0.03, 1.34] |
| Shovels | 0.28 (0.18) | 0.22 (0.12) | 0.60 | 0.33 [-0.26, 0.95] |
| Pens | 0.12 (0.15) | 0.13 (0.16) | 0.33 | -0.05 [-0.63, 0.54] |
| Corridors | 0.48 (0.15) | 0.42 (0.20) | 0.45 | 0.25 [-0.32, 0.85] |
| Houses | 0.47 (0.16) | 0.39 (0.18) | 0.77 | 0.39 [-0.19, 1.03] |
| Words | 0.33 (0.16) | 0.33 (0.20) | 0.32 | -0.01 [-0.58, 0.57] |
| Pseudowords | 0.33 (0.23) | 0.32 (0.21) | 0.32 | 0.02 [-0.56, 0.60] |

*Right LOTC:*

| Category | Late Learners M (SD) | Non-signers M (SD) | $BF_{10}$ | δ [95% CrI] |
|---|---|---|---|---|
| Adult Faces | 0.39 (0.14) | 0.40 (0.18) | 0.32 | -0.03 [-0.61, 0.53] |
| Child Faces | 0.34 (0.12) | 0.38 (0.18) | 0.39 | -0.18 [-0.76, 0.39] |
| Bodies | 0.57 (0.19) | 0.50 (0.13) | 0.70 | 0.37 [-0.20, 1.02] |
| Hands | 0.43 (0.17) | 0.31 (0.17) | 1.70 | 0.57 [-0.04, 1.23] |
| Shovels | 0.20 (0.12) | 0.21 (0.10) | 0.35 | -0.11 [-0.70, 0.45] |
| Pens | 0.14 (0.15) | 0.17 (0.19) | 0.36 | -0.14 [-0.74, 0.45] |
| Corridors | 0.56 (0.15) | 0.46 (0.20) | 0.88 | 0.43 [-0.15, 1.07] |
| Houses | 0.53 (0.13) | 0.45 (0.12) | 1.25 | 0.50 [-0.10, 1.17] |
| Words | 0.18 (0.13) | 0.20 (0.16) | 0.34 | -0.09 [-0.68, 0.48] |
| Pseudowords | 0.11 (0.21) | 0.15 (0.15) | 0.41 | -0.20 [-0.81, 0.37] |

*Left lateral VTC:*

| Category | Late Learners M (SD) | Non-signers M (SD) | $BF_{10}$ | δ [95% CrI] |
|---|---|---|---|---|
| Adult Faces | 0.67 (0.16) | 0.58 (0.22) | 0.66 | 0.35 [-0.23, 0.97] |
| Child Faces | 0.64 (0.14) | 0.55 (0.21) | 0.74 | 0.38 [-0.19, 1.01] |
| Bodies | 0.40 (0.15) | 0.34 (0.18) | 0.52 | 0.28 [-0.28, 0.92] |
| Hands | 0.44 (0.18) | 0.28 (0.15) | 7.10 | 0.82 [0.15, 1.52] |
| Shovels | 0.39 (0.16) | 0.34 (0.14) | 0.46 | 0.24 [-0.34, 0.85] |
| Pens | 0.22 (0.11) | 0.18 (0.13) | 0.42 | 0.21 [-0.37, 0.82] |
| Corridors | 0.58 (0.16) | 0.48 (0.16) | 1.40 | 0.53 [-0.07, 1.19] |
| Houses | 0.59 (0.18) | 0.48 (0.14) | 1.41 | 0.53 [-0.08, 1.19] |
| Words | 0.53 (0.16) | 0.60 (0.24) | 0.50 | -0.27 [-0.89, 0.31] |
| Pseudowords | 0.53 (0.13) | 0.62 (0.18) | 1.03 | -0.46 [-1.12, 0.13] |

*Right lateral VTC:*

| Category | Late Learners M (SD) | Non-signers M (SD) | $BF_{10}$ | δ [95% CrI] |
|---|---|---|---|---|
| Adult Faces | 0.78 (0.10) | 0.72 (0.22) | 0.48 | 0.25 [-0.31, 0.86] |
| Child Faces | 0.74 (0.12) | 0.68 (0.19) | 0.50 | 0.26 [-0.29, 0.88] |
| Bodies | 0.52 (0.15) | 0.48 (0.19) | 0.37 | 0.15 [-0.40, 0.78] |
| Hands | 0.38 (0.19) | 0.33 (0.18) | 0.42 | 0.21 [-0.35, 0.81] |
| Shovels | 0.30 (0.14) | 0.34 (0.10) | 0.46 | -0.25 [-0.86, 0.32] |
| Pens | 0.26 (0.16) | 0.24 (0.20) | 0.34 | 0.09 [-0.48, 0.69] |
| Corridors | 0.57 (0.15) | 0.50 (0.16) | 0.63 | 0.35 [-0.23, 0.97] |
| Houses | 0.54 (0.17) | 0.51 (0.13) | 0.38 | 0.16 [-0.41, 0.75] |
| Words | 0.33 (0.17) | 0.36 (0.18) | 0.37 | -0.14 [-0.74, 0.43] |
| Pseudowords | 0.35 (0.25) | 0.35 (0.18) | 0.32 | 0.02 [-0.56, 0.60] |

*Left medial VTC:*

| Category | Late Learners M (SD) | Non-signers M (SD) | $BF_{10}$ | δ [95% CrI] |
|---|---|---|---|---|
| Adult Faces | 0.45 (0.12) | 0.34 (0.16) | 2.06 | 0.60 [-0.02, 1.25] |
| Child Faces | 0.32 (0.16) | 0.22 (0.15) | 1.32 | 0.51 [-0.08, 1.16] |
| Bodies | 0.19 (0.11) | 0.13 (0.16) | 0.56 | 0.31 [-0.25, 0.95] |
| Hands | 0.01 (0.11) | 0.04 (0.12) | 0.46 | -0.24 [-0.83, 0.32] |
| Shovels | 0.17 (0.11) | 0.13 (0.14) | 0.54 | 0.29 [-0.29, 0.91] |
| Pens | 0.00 (0.06) | 0.05 (0.09) | 0.87 | -0.42 [-1.07, 0.18] |
| Corridors | 0.63 (0.15) | 0.60 (0.18) | 0.37 | 0.16 [-0.41, 0.75] |
| Houses | 0.53 (0.16) | 0.45 (0.15) | 0.73 | 0.38 [-0.21, 1.02] |
| Words | 0.31 (0.14) | 0.25 (0.18) | 0.51 | 0.28 [-0.28, 0.91] |
| Pseudowords | 0.23 (0.15) | 0.22 (0.15) | 0.33 | 0.05 [-0.53, 0.64] |

*Right medial VTC:*

| Category | Late Learners M (SD) | Non-signers M (SD) | $BF_{10}$ | δ [95% CrI] |
|---|---|---|---|---|
| Adult Faces | 0.40 (0.18) | 0.46 (0.22) | 0.42 | -0.21 [-0.81, 0.34] |
| Child Faces | 0.30 (0.19) | 0.36 (0.22) | 0.43 | -0.22 [-0.81, 0.35] |
| Bodies | 0.28 (0.17) | 0.16 (0.16) | 1.78 | 0.57 [-0.03, 1.26] |
| Hands | 0.03 (0.11) | 0.03 (0.12) | 0.32 | -0.02 [-0.58, 0.56] |
| Shovels | 0.14 (0.14) | 0.16 (0.11) | 0.35 | -0.11 [-0.70, 0.45] |
| Pens | 0.00 (0.11) | 0.03 (0.12) | 0.38 | -0.16 [-0.76, 0.42] |
| Corridors | 0.68 (0.14) | 0.66 (0.20) | 0.35 | 0.11 [-0.45, 0.70] |
| Houses | 0.57 (0.14) | 0.53 (0.20) | 0.38 | 0.17 [-0.40, 0.77] |
| Words | 0.20 (0.17) | 0.22 (0.17) | 0.34 | -0.09 [-0.68, 0.49] |
| Pseudowords | 0.07 (0.11) | 0.10 (0.15) | 0.40 | -0.19 [-0.80, 0.38] |

**Supplemental Table 5. Bayesian independent-samples t-tests for Distinctiveness of late learning signers vs non-signers for all anatomical ROIs**

N late learning signers = 16, N non-signers = 20. $BF_{10}$ = bayes factor in favor of the alternative hypothesis; δ = posterior standardized effect size; CrI = credible interval. Positive δ values indicate higher values for signers compared to non-signers. Bayes factors were calculated using a cauchy prior with r = 0.707.

*Left LOTC:*

| Category | Late Learners M (SD) | Non-signers M (SD) | $BF_{10}$ | δ [95% CrI] |
|---|---|---|---|---|
| Adult Faces | 2.19 (2.04) | 1.95 (1.81) | 0.34 | 0.09 [-0.48, 0.66] |
| Child Faces | 1.67 (2.25) | 1.05 (1.10) | 0.51 | 0.27 [-0.28, 0.89] |
| Bodies | 7.71 (3.77) | 7.87 (4.29) | 0.32 | -0.03 [-0.60, 0.56] |
| Hands | 13.21 (5.43) | 8.80 (5.83) | 2.46 | 0.64 [0.01, 1.31] |
| Shovels | 2.81 (3.15) | 1.83 (1.91) | 0.54 | 0.29 [-0.29, 0.91] |
| Pens | 1.28 (1.79) | 0.74 (0.92) | 0.55 | 0.30 [-0.27, 0.93] |
| Corridors | 2.56 (2.72) | 3.58 (2.92) | 0.51 | -0.27 [-0.89, 0.30] |
| Houses | 1.60 (2.05) | 1.16 (1.62) | 0.39 | 0.18 [-0.38, 0.78] |
| Words | 6.52 (4.33) | 8.80 (7.30) | 0.52 | -0.28 [-0.90, 0.30] |
| Pseudowords | 7.75 (5.15) | 7.93 (6.01) | 0.32 | -0.02 [-0.61, 0.55] |

*Right LOTC:*

| Category | Late Learners M (SD) | Non-signers M (SD) | $BF_{10}$ | δ [95% CrI] |
|---|---|---|---|---|
| Adult Faces | 3.18 (2.81) | 3.96 (3.71) | 0.39 | -0.18 [-0.77, 0.37] |
| Child Faces | 2.25 (1.67) | 1.85 (1.46) | 0.41 | 0.19 [-0.36, 0.79] |
| Bodies | 12.54 (5.63) | 14.86 (5.73) | 0.57 | -0.31 [-0.94, 0.27] |
| Hands | 12.03 (4.69) | 8.82 (5.92) | 1.06 | 0.47 [-0.12, 1.11] |
| Shovels | 0.84 (0.87) | 1.14 (1.43) | 0.40 | -0.19 [-0.80, 0.37] |
| Pens | 0.78 (1.24) | 0.51 (0.61) | 0.43 | 0.22 [-0.36, 0.84] |
| Corridors | 4.20 (4.09) | 3.53 (3.16) | 0.37 | 0.14 [-0.42, 0.73] |
| Houses | 1.99 (2.35) | 1.30 (1.41) | 0.51 | 0.28 [-0.29, 0.90] |
| Words | 2.28 (1.94) | 5.44 (5.15) | 2.42 | -0.63 [-1.32, 0.01] |
| Pseudowords | 2.25 (1.75) | 4.78 (5.37) | 1.12 | -0.48 [-1.14, 0.11] |

*Left lateral VTC:*

| Category | Late Learners M (SD) | Non-signers M (SD) | $BF_{10}$ | δ [95% CrI] |
|---|---|---|---|---|
| Adult Faces | 9.95 (6.37) | 7.50 (7.07) | 0.51 | 0.27 [-0.29, 0.88] |
| Child Faces | 5.74 (3.93) | 4.79 (4.76) | 0.38 | 0.16 [-0.39, 0.75] |
| Bodies | 9.40 (6.92) | 6.50 (5.84) | 0.66 | 0.35 [-0.21, 1.00] |
| Hands | 10.02 (9.01) | 4.29 (3.68) | 3.92 | 0.72 [0.07, 1.40] |
| Shovels | 2.84 (2.77) | 2.33 (3.55) | 0.35 | 0.12 [-0.46, 0.70] |
| Pens | 0.83 (1.34) | 0.39 (0.64) | 0.62 | 0.33 [-0.25, 0.97] |
| Corridors | 0.27 (0.59) | 0.38 (0.54) | 0.37 | -0.14 [-0.73, 0.43] |
| Houses | 0.29 (0.34) | 0.14 (0.31) | 0.68 | 0.36 [-0.22, 1.00] |
| Words | 24.06 (16.02) | 23.80 (14.30) | 0.32 | 0.02 [-0.56, 0.60] |
| Pseudowords | 21.50 (11.75) | 21.40 (14.65) | 0.32 | 0.01 [-0.57, 0.59] |

*Right lateral VTC:*

| Category | Late Learners M (SD) | Non-signers M (SD) | $BF_{10}$ | δ [95% CrI] |
|---|---|---|---|---|
| Adult Faces | 14.56 (5.28) | 14.00 (8.81) | 0.33 | 0.05 [-0.52, 0.62] |
| Child Faces | 9.41 (3.75) | 9.79 (6.40) | 0.33 | -0.06 [-0.62, 0.51] |
| Bodies | 18.66 (11.18) | 17.34 (9.12) | 0.34 | 0.10 [-0.46, 0.71] |
| Hands | 7.63 (5.47) | 6.60 (8.31) | 0.35 | 0.11 [-0.45, 0.69] |
| Shovels | 0.77 (1.13) | 0.79 (1.50) | 0.32 | -0.01 [-0.59, 0.56] |
| Pens | 0.42 (0.96) | 0.43 (1.03) | 0.32 | -0.01 [-0.59, 0.58] |
| Corridors | 0.72 (0.77) | 1.20 (1.55) | 0.53 | -0.29 [-0.91, 0.29] |
| Houses | 0.62 (0.81) | 0.66 (1.15) | 0.33 | -0.03 [-0.61, 0.54] |
| Words | 9.53 (8.29) | 10.92 (9.30) | 0.35 | -0.11 [-0.71, 0.46] |
| Pseudowords | 8.65 (7.31) | 7.54 (7.15) | 0.35 | 0.12 [-0.46, 0.71] |

*Left medial VTC:*

| Category | Late Learners M (SD) | Non-signers M (SD) | $BF_{10}$ | δ [95% CrI] |
|---|---|---|---|---|
| Adult Faces | 0.95 (1.37) | 0.62 (1.37) | 0.39 | 0.17 [-0.39, 0.76] |
| Child Faces | 0.57 (0.86) | 0.35 (0.76) | 0.42 | 0.21 [-0.34, 0.81] |
| Bodies | 0.85 (1.09) | 0.38 (1.21) | 0.56 | 0.31 [-0.25, 0.94] |
| Hands | 0.28 (0.55) | 0.45 (1.05) | 0.37 | -0.15 [-0.73, 0.41] |
| Shovels | 1.30 (1.95) | 0.63 (1.12) | 0.62 | 0.33 [-0.25, 0.96] |
| Pens | 0.13 (0.24) | 0.27 (0.62) | 0.42 | -0.22 [-0.82, 0.37] |
| Corridors | 12.53 (7.04) | 14.19 (8.23) | 0.38 | -0.16 [-0.75, 0.41] |
| Houses | 7.23 (5.07) | 9.13 (7.01) | 0.45 | -0.23 [-0.85, 0.33] |
| Words | 6.42 (4.63) | 2.99 (2.57) | 5.93 | 0.80 [0.14, 1.52] |
| Pseudowords | 5.64 (3.02) | 3.31 (3.09) | 2.23 | 0.62 [-0.00, 1.31] |

*Right medial VTC:*

| Category | Late Learners M (SD) | Non-signers M (SD) | $BF_{10}$ | δ [95% CrI] |
|---|---|---|---|---|
| Adult Faces | 0.64 (0.99) | 2.48 (4.24) | 0.97 | -0.45 [-1.09, 0.13] |
| Child Faces | 0.49 (0.99) | 1.33 (2.20) | 0.69 | -0.37 [-0.99, 0.21] |
| Bodies | 2.01 (2.39) | 2.14 (3.02) | 0.33 | -0.03 [-0.61, 0.56] |
| Hands | 0.40 (1.03) | 0.45 (1.37) | 0.32 | -0.03 [-0.60, 0.54] |
| Shovels | 1.14 (1.85) | 0.53 (0.78) | 0.64 | 0.34 [-0.24, 0.97] |
| Pens | 0.08 (0.21) | 0.06 (0.12) | 0.34 | 0.10 [-0.48, 0.70] |
| Corridors | 18.83 (7.50) | 18.23 (13.01) | 0.33 | 0.04 [-0.52, 0.62] |
| Houses | 13.18 (8.90) | 12.40 (10.30) | 0.33 | 0.06 [-0.51, 0.64] |
| Words | 3.02 (2.86) | 3.07 (2.83) | 0.32 | -0.01 [-0.59, 0.57] |
| Pseudowords | 2.68 (3.40) | 2.31 (3.01) | 0.34 | 0.09 [-0.48, 0.68] |

**Supplemental Table 6. Bayesian independent-samples t-tests for Nr. of selective voxel analysis of late learning signers vs non-signers for all anatomical ROIs**

N late learning signers = 16, N non-signers = 20. $BF_{10}$ = bayes factor in favor of the alternative hypothesis; δ = posterior standardized effect size; CrI = credible interval. Positive δ values indicate higher values for signers compared to non-signers. Bayes factors were calculated using a cauchy prior with r = 0.707.

| Category | Signers M (SD) | Non-signers M (SD) | $BF_{10}$ | δ [95% CrI] |
|---|---|---|---|---|
| Overall | 85.19 (8.89) | 88.56 (5.68) | 0.69 | -0.36 [-0.95, 0.18] |
| Adult Faces | 86.11 (10.12) | 92.22 (11.46) | 1.07 | -0.46 [-1.06, 0.10] |
| Child Faces | 91.39 (8.16) | 90.83 (8.70) | 0.31 | 0.05 [-0.48, 0.62] |
| Bodies | 79.17 (15.28) | 87.50 (11.94) | 1.30 | -0.49 [-1.09, 0.08] |
| Hands | 89.17 (15.56) | 87.78 (11.34) | 0.32 | 0.07 [-0.47, 0.63] |
| Shovels | 87.22 (10.83) | 91.11 (9.94) | 0.54 | -0.29 [-0.89, 0.27] |
| Pens | 80.56 (14.25) | 85.83 (10.74) | 0.62 | -0.33 [-0.92, 0.23] |
| Corridors | 87.78 (13.92) | 87.78 (10.60) | 0.31 | -0.00 [-0.54, 0.55] |
| Houses | 81.94 (16.61) | 86.67 (8.34) | 0.51 | -0.28 [-0.87, 0.28] |
| Words | 82.78 (13.23) | 86.39 (11.75) | 0.43 | -0.22 [-0.81, 0.32] |
| Pseudowords | 85.83 (12.42) | 89.44 (11.67) | 0.44 | -0.23 [-0.81, 0.31] |

**Supplemental Table 7. Bayesian independent-samples t-tests on the accuracy in the oddball-task between signers and non-signers**

N of both groups = 20. $BF_{10}$ = bayes factor in favor of the alternative hypothesis; δ = posterior standardized effect size; CrI = credible interval. Positive δ values indicate higher values for signers compared to non-signers. Bayes factors were calculated using a cauchy prior with r = 0.707.

| ROI | Signers M (SD) | Non-signers M (SD) | BF10 | δ [95% CrI] |
|---|---|---|---|---|
| Left LOTC | 6517.80 (1136.27) | 6625.15 (1233.31) | 0.32 | -0.07 [-0.63, 0.48] |
| Right LOTC | 7139.85 (327.79) | 7457.50 (873.99) | 0.41 | -0.20 [-0.78, 0.34] |
| Left lateral VTC | 1716.50 (427.80) | 2146.50 (489.76) | 8.12 | -0.79 [-1.46-0.17] |
| Right lateral VTC | 1959.95 (631.53) | 1995.55 (605.28) | 0.31 | -0.04 [-0.60, 0.50] |
| Left medial VTC | 1790.00 (441.93) | 1911.60 (448.27) | 0.42 | -0.21 [-0.79, 0.33] |
| Right medial VTC | 1849.50 (398.65) | 1841.25 (426.60) | 0.31 | 0.02 [-0.53, 0.57] |

**Supplemental Table 8. Bayesian independent-samples t-tests of total nr. of voxels of anatomical ROIs between signers and non-signers**

N of both groups = 20. $BF_{10}$ = bayes factor in favor of the alternative hypothesis; δ = posterior standardized effect size; CrI = credible interval. Positive δ values indicate higher values for signers compared to non-signers. Bayes factors were calculated using a cauchy prior with r = 0.707.
